# GPBench: A benchmark framework for genomic prediction

**DOI:** 10.64898/2026.10.31.762999

**Authors:** Xiaowei Zhang, Hanwen Luo, Geng Li, Xuerui Zhang, Siying Wang, Liliang Liu, Maozu Guo, Qiaoyu Tan, Carlotta Domeniconi, Jian Chen, Jun Wang, Guoxian Yu

## Abstract

Genomic prediction (GP) has emerged as a powerful data mining strategy for disease risk assessment and molecular breeding. GP leverages genome-wide genetic variations and phenotypic information through a computational model to estimate genetic potential or disease risk. However, the field of GP still faces several major challenges: (i) dataset bias and insufficient representativeness; (ii) the lack of standardized evaluation protocols and a systematic benchmarking framework; (iii) heavy reliance on empirical expertise for experimental result interpretation, which hinders large-scale and systematic analysis. These issues make it difficult to conduct systematic cross-species evaluation and comparison of GP methods, and their authentic applications. To address these challenges, we established a standardized and extensible GP benchmark framework—GPBench. We first collected and harmonized public datasets from **16 species**, covering animals, plants, and microorganisms, comprising **57 traits, 40**,**821 samples, and 493**,**821 single-nucleotide polymorphisms**. Then, using the standardized datasets ^1^, the framework evaluates 21 GP models^2^ and provides a unified assessment across four dimensions: effectiveness, sensitivity, robustness, and scalability. Furthermore, we developed a large language model-based intelligent analysis system to facilitate the interpretation of GP experiments and model recommendations. GPBench provides the community with a fair, unified, and scalable platform (www.sdu-idea.cn/GPBench/) and an open-source library for comparing and optimizing GP methods, ultimately accelerating the application of GP in life sciences and biomedical research.

## 1 Introduction

Genomic prediction (GP) is a data mining approach that integrates genome-wide single-nucleotide polymorphism (SNP) markers with phenotypic data to predict the genetic potential of individuals for specific target traits (i.e., color, height, and cancer) [17]. In human health research, GP has been further extended to disease risk scoring [5], where it is commonly formulated as a classification task to quantify an individual’s genetic susceptibility to complex diseases, including cancer [25], psychiatric disorders [2], and age-related macular degeneration [5, 57]. In agriculture and animal husbandry, GP is also known as genomic selection (GS) and has been widely applied to the early selection of complex traits, primarily as a regression task. For example, it can be used to estimate crop yield [9] and livestock milk production [42]. This thereby reduces the need of extensive phenotypic evaluations and accelerating genetic improvement research in plants and animals.

With the advancement of sequencing technologies, GP methods have evolved rapidly in recent years. Traditional statistical genetics models, such as RR-BLUP [10] and GBLUP [13], remain widely used due to their robustness, efficiency, and biological interpretability, and are commonly adopted as canonical baselines. In parallel, machine learning (ML) and deep learning (DL) approaches—including random forests [4] and neural networks [29]—have gained increasing attention for their ability to capture nonlinear effects and highorder interactions among SNP markers. Beyond purely data-driven modeling, integrating biological knowledge, such as biological pathways [3], transcriptomics [18], epigenomics [19], and metabolomics [43], has emerged as a promising direction for improving and understanding the genetic architecture of complex traits.

Despite this rapid methodological progress, the GP field still lacks a standardized and systematic benchmarking framework for evaluating and comparing models in a fair and reproducible manner. Several critical challenges remain unresolved. **First**, *dataset bias and insufficient representativeness* continue to hinder reliable assessment [60]. Most existing studies focus on a single species or a small number of traits, and the absence of benchmarks spanning diverse biological domains limits the ability to assess cross-species generalization. **Second**, *the lack of standardized evaluation protocols* complicates meaningful comparison across studies. Current evaluations predominantly emphasize prediction accuracy or error-based metrics, while largely neglecting other important dimensions such as sensitivity to data scale, robustness to noise or missingness, and computational scalability. Consequently, reported performance gains are often incomplete or difficult to interpret in real-world settings. **Third**, *the interpretation of experimental results is highly experience-driven and difficult to scale*. As GP evaluations grow in size, covering many datasets, models, and experimental conditions, manual analysis becomes increasingly subjective and inefficient.

To address these challenges, we develop a standardized, extensible, and generalizable GP benchmarking framework—GPBench. Our main contributions are summarized as follows: **(1) Standardized and systematically curated multi-species datasets**. We collected multiple publicly available genotype–phenotype datasets covering animals, plants, and microorganisms, encompassing 16 species and 57 traits, thereby alleviating issues of dataset bias and limited representativeness. **(2) Unified and scalable GP evaluation framework**. GPBench systematically benchmarks 21 widely-used GP algorithms across four complementary dimensions: effectiveness, sensitivity, robustness, and scalability. The framework supports fair comparisons across multi-species and multi-trait scenarios, while remaining easily extensible to new datasets and methods. **(3) Intelligent and automated data analysis platform**. We release an open-source GPBench library and an online platform GPBench that integrates an intelligent data analysis module based on large language models (LLMs). This module supports automated result summarization and evidence-based algorithm recommendation for previously unexplored application scenarios, substantially reducing reliance on manual, experience-driven interpretation. **(4) Empirical insights from large-scale benchmarking**. Our results show that statistical genetics–based methods remain competitive and stable in effectiveness and sensitivity across most species, serving as strong baselines. Feature engineering–based methods, particularly tree-based models, achieve top performance across multiple evaluation dimensions in several species. In contrast, sequence-based deep learning models exhibit substantial variability, with robustness and generalization performance remaining limited under realistic perturbations.

## 2 Related Work

To understand the current state of development in GP methods, we review their evolution from 2001 to 2025. Based on their modeling assumptions and representation strategies, we broadly categorize existing GP methods into three groups: (1) statistical genetics–based methods, (2) feature-engineered classical machine learning methods, and (3) sequence representation–based deep learning methods.

### 2.1 Statistical genetics-based methods

Statistical genetics-based methods typically require a prior assumption about the genetic model structure, often modeling the phenotype as a combination of additive effects from multiple SNPs. The basic form can be written as:

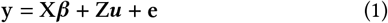

where y is the phenotype vector, X and Z are the associated design matrices, ***β*** and ***u*** are vectors of fixed and random effects, and e is a random residual vector. Typical methods include the best linear unbiased prediction (BLUP) and Bayesian alphabet methods (i.e., BayesA [31], BayesB [31], BayesC*π* [14]). For example, genomic BLUP (GBLUP) estimates genomic breeding values by integrating the whole-genome information, using a genomic relationship matrix calculated from SNP markers to capture genetic similarities between individuals [13]. The Bayesian alphabet extends the statistical genetic framework by introducing prior assumptions on SNP effect distributions. Although these methods are computationally efficient and highly interpretable, they struggle to capture complex nonlinear genetic effects and higher-order interactions, making them inadequate for modeling complex genetic relationships.

### 2.2 Feature-engineered classical ML methods

Feature-engineered classical machine learning methods formulate the genomic prediction task as a supervised learning problem that maps high-dimensional genotypic feature spaces to continuous phenotypic outcomes, aiming to accurately predict individual traits by mining complex relationships between genotypes and phenotypes. Specifically, these approaches typically take informative SNPs as input features and individual samples as training instances, and iteratively optimize model parameters to achieve robust performance on both training and test sets. Classical methods include kernel-based methods (SVR [28]), tree-based ensemble models (random forest (RF) [4], XGboost [7], and LightGBM [54]). Such models surpass traditional statistical methods in capturing nonlinear patterns and higher-order interactions through flexible nonparametric modeling and effective feature selection. Furthermore, automated machine learning (AutoML) [15] lowers the technical barriers to hyperparameter tuning, thereby accelerating the application of machine learning models in disease risk assessment and plant and animal breeding.

### 2.3 Sequence representation-based DL methods

Sequence representation-based deep learning methods show strong potential by automatically learning complex feature representations through multi-layer nonlinear networks. Currently, the main architectures used in GP can be broadly categorized into convolutional neural network (CNN)-based and hybrid architecture-based models. **(1) CNN-based models**, such as DeepGS [29], G2PDeep [59], and DNNGP [48], exploit their local and hierarchical receptive fields to capture sequence patterns and identify key SNP features. (2) **Hybrid architecture-based models** integrate the strengths of multiple model structures by combining CNNs with long short-term memory (LSTM) networks or attention mechanisms. This design allows the models to simultaneously capture local sequence features and global contextual information, thereby enhancing their ability to model complex genomic regulatory patterns or genotype–phenotype associations, e.g. GPformer [51], DeepCCR [30], and Cropformer [47]. In addition, LLMs can learn contextual dependencies from large-scale genomic data, offering new opportunities for genomic prediction [32]. These methods offer strong nonlinear modeling and feature representation capabilities on large-scale genomic data, but need substantial data and computational resources, facing challenges in interpretability and generalization.

## 3 Genomic Prediction Benchmark

To enable a systematic evaluation of GP algorithms, we construct an evaluation framework that spans data, algorithms, and evaluation metrics, as shown in Figure 2.

**Figure 1.**
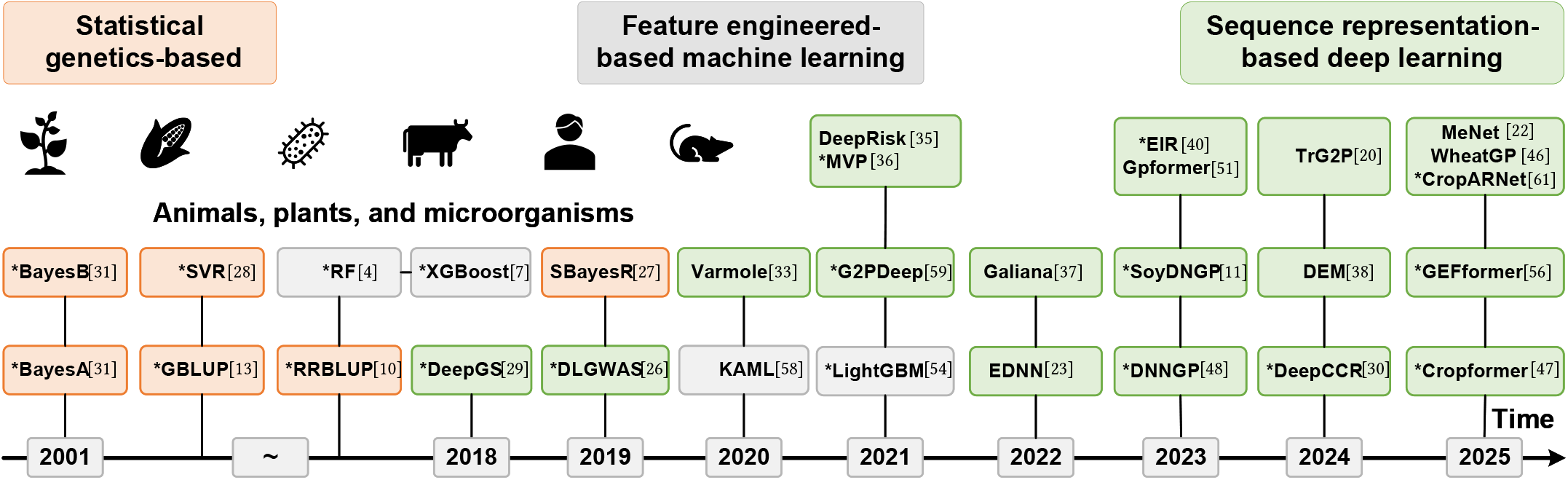
The evolution of genomic prediction: from traditional statistical models to AI-driven paradigm shifts.

**Figure 2.**
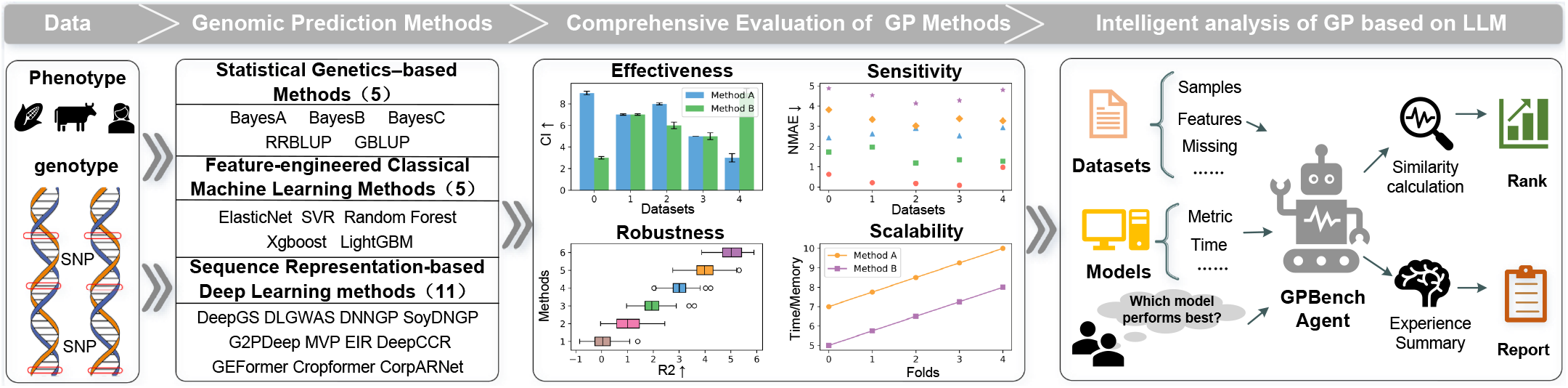
The GPBench framework. Using the benchmark datasets, we quantitatively compare 21 mainstream GP methods in terms of effectiveness, sensitivity, robustness, and scalability. Furthermore, we develop an agent for the GP community to automatically generate evaluation reports and recommend suitable methods.

### 3.1 Benchmark Datasets

To comprehensively evaluate algorithmic performance and systematically analyze the limitations of existing GP methods, we collected a diverse set of widely used datasets spanning 16 species (Figure 3). Through systematic quality control, data preprocessing, and genotype encoding, we constructed a unified benchmark dataset (see Appendix A.1).

**Figure 3.**
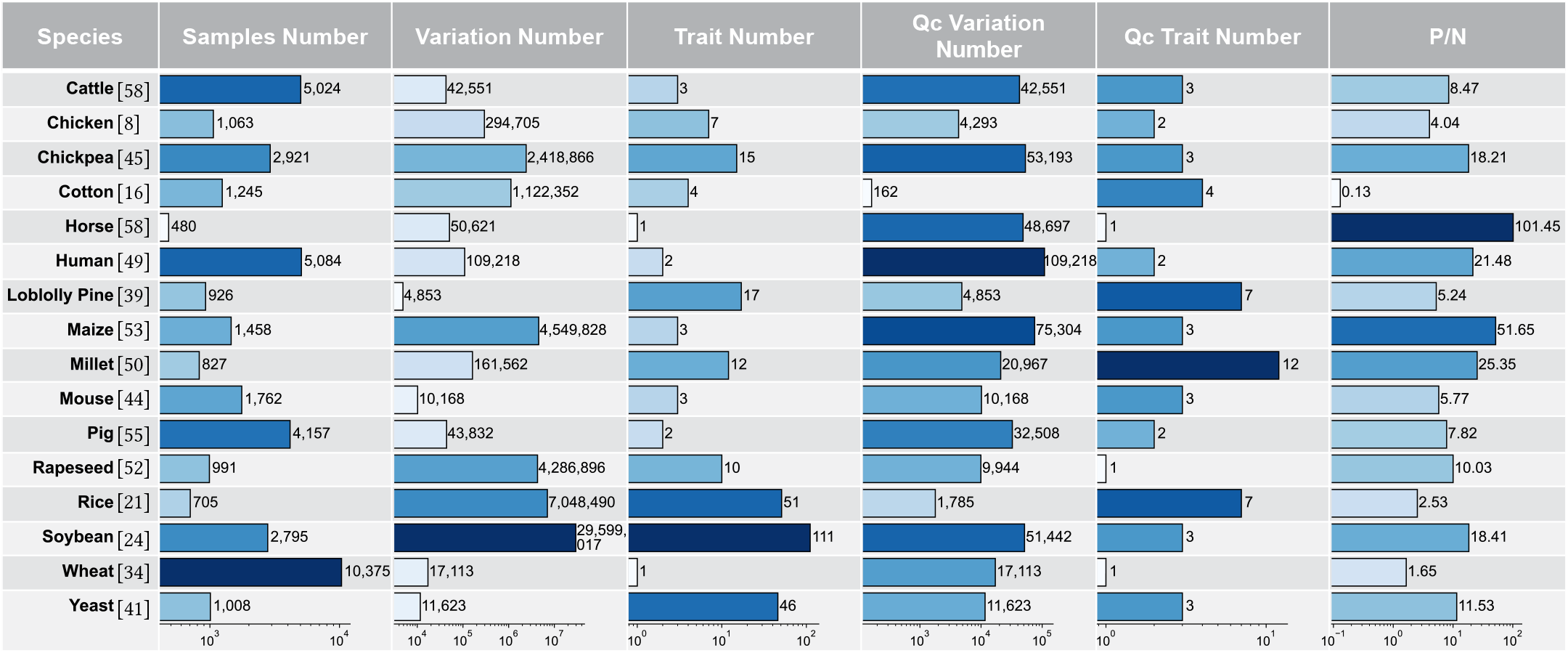
Characteristics of the benchmark datasets, including the number of samples, variants, traits, and the feature-to-sample ratio (*P*/*N*) of each species.

The resulting benchmark dataset exhibits three prominent characteristics. **(i) Broad species coverage**. The datasets span a wide range of model organisms, including animals, plants, and microorganisms, encompassing diverse genetic backgrounds and breeding objectives. Such species diversity enables a comprehensive evaluation of GP methods across heterogeneous biological scenarios. **(ii) Rich and diverse phenotypic data**. A total of 57 traits were collected and analyzed using kernel density estimation (see Supplementary Fig. S1). These traits have diverse phenotypic distribution patterns. Some traits approximately follow unimodal normal distributions (e.g., plant height and grain weight), whereas others exhibit pronounced multimodal or skewed non-normal distributions (e.g., grain length and flowering time). This diversity in phenotypic distributions more realistically reflects the complexity of genetic studies and real-world application scenarios. **(iii) Diverse and representative feature spaces**. The datasets cover feature spaces ranging from relatively low-dimensional to ultra-high-dimensional. Low-dimensional datasets contain a limited number of molecular markers, whereas high-dimensional datasets typically include tens of thousands of SNPs. Overall, most datasets exhibit a ‘high-dimensional, small-sample-size’ characteristic, which is one of the core genomic prediction challenges.

### 3.2 Benchmark Algorithms

To comprehensively evaluate the performance, we selected 21 state-of-the-art methods as core benchmark algorithms, covering all four major paradigms. Statistical genetics-based methods include rrBLUP [10], GBLUP [13], BayesA [31], BayesB [31], and BayesC [14]; Feature-engineered classical machine learning methods include ElasticNet [62], SVR [28], RF [4], XGBoost [7], and LightGBM [54]; CNN-based methods include DeepGS [29], DLGWAS [26], G2PDeep [59], MVP [36], and DNNGP [48]; and Hybrid architecture-based methods include EIR [40], SoyDNGP [11], DeepCCR [30], Cropformer [47], GEFormer [56], and CropARNet [61]. More details about these benchmark algorithms are deferred to Appendix A.2. We adopted a standardized workflow to assemble 21 algorithms into a unified evaluation framework (see Appendix A.4.2). We adhered to the original model designs and code logic to ensure the reproducibility and comparability of experimental results, thereby providing a reliable and robust evaluation standard for GP research.

### 3.3 Benchmark Evaluations

We construct a comprehensive evaluation framework to rigorously assess benchmark algorithms along four key dimensions:

**(i) Effectiveness** is a canonically used evaluation dimension in GP studies and is typically assessed using regression metrics such as the Pearson correlation coefficient (PCC) and mean absolute error (MAE), which are insufficient to fully characterize algorithmic effectiveness. To address this issue, we first adopted a broader set of evaluation metrics, including the consistent index (CI) [51], root mean square error (RMSE), and the coefficient of determination (R2), to comprehensively assess the performance of existing methods. In addition, we further investigated the impact of genetic structure differences on model predictive performance by conducting comparative experiments on datasets with diverse genetic backgrounds and population structures, thereby systematically analyzing the behavior of different methods under genetic heterogeneity.

**(ii) Sensitivity** is defined as the response of GP model performance to changes in data features, sample size, and random initialization in this paper. Specifically, we assess the sensitivity of GP methods by evaluating them under different random seed settings and across datasets of varying scales (in terms of sample size and feature dimensionality), and by quantifying performance fluctuations and standard deviations under cross-validation, thereby revealing how existing models respond to external variations.

**(3) Robustness** is primarily used to assess the performance of GP algorithms under perturbations such as biological sequencing noise and missing values, which represent key challenges in current GP research. We posit that a robust GP algorithm should maintain high predictive accuracy and strong fitting capability in noisy environments, exhibit strong resistance to data perturbations, and have limited performance degradation within an acceptable range.

**(4) Scalability** evaluates the computational capability of GP algorithms when processing large-scale genotype data, with a focus on time and space complexity. Algorithms with good scalability should be able to run efficiently on datasets of different sizes, avoiding significant performance degradation or memory overflow issues. An efficient algorithm should minimize computational resource consumption while achieving fast convergence.

### 3.4 Intelligent data analysis

Considering that the interpretation of experimental results across multiple datasets is highly experience-dependent and difficult to scale, we develop an LLM-based agent (supported by Qwen-plus) for GP result analysis and algorithm recommendation [12], assisting users in selecting appropriate genomic prediction methods under different data characteristics and application scenarios. This module supports two types of tasks.

#### (i) Result analysis task

When users are only interested in analyzing existing experimental results, they can directly query Agent in natural language, for example: “What patterns do the models exhibit for the maize DTT (Days to tasseling) trait, and what insights can be derived?” The Agent retrieves and reasons over prior algorithm performance records and returns corresponding analytical conclusions (see Appendix B.4, Fig.S9).

### (ii) Algorithm recommendation task

When users provide a new dataset and request a GP algorithm recommendation (see Appendix B.4, Fig.S10), we formulate the problem as a context-aware, experience-driven decision process. The new dataset X_*d*_ is depicted as

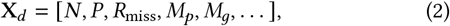

where *N* denotes the number of samples, *P* is the feature dimensionality (number of SNPs), *R*_miss_ is the missing rate, *M*_*p*_ and *M*_*g*_ describe the mean values of phenotype and genotype.

Based on this representation, Agent retrieves similar historical scenarios and integrates the multi-dimensional performance of candidate algorithms under user preference constraints to produce a ranked list *π* of GP algorithm recommendations:

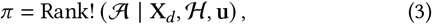

where denotes the set of algorithms, denotes accumulated historical knowledge, and u represents user preferences. The resulting ranking is accompanied by interpretable justifications to support algorithm selection under diverse data conditions.

## 4 Results

### 4.1 Effectiveness analysis

#### 4.1.1 Analysis of multi-species experiment

We conducted a systematic evaluation of 21 methods, as summarized in Table 1. For regression tasks, performance was assessed using PCC, CI [51], MAE (see Appendix B.1.1, Table S4), RMSE (Table S5), and *R*^2^ (Table S6); for classification tasks, Accuracy, Precision, Recall, F1-score were used (see Appendix B.1.2, Fig. S2).

**Table 1.**
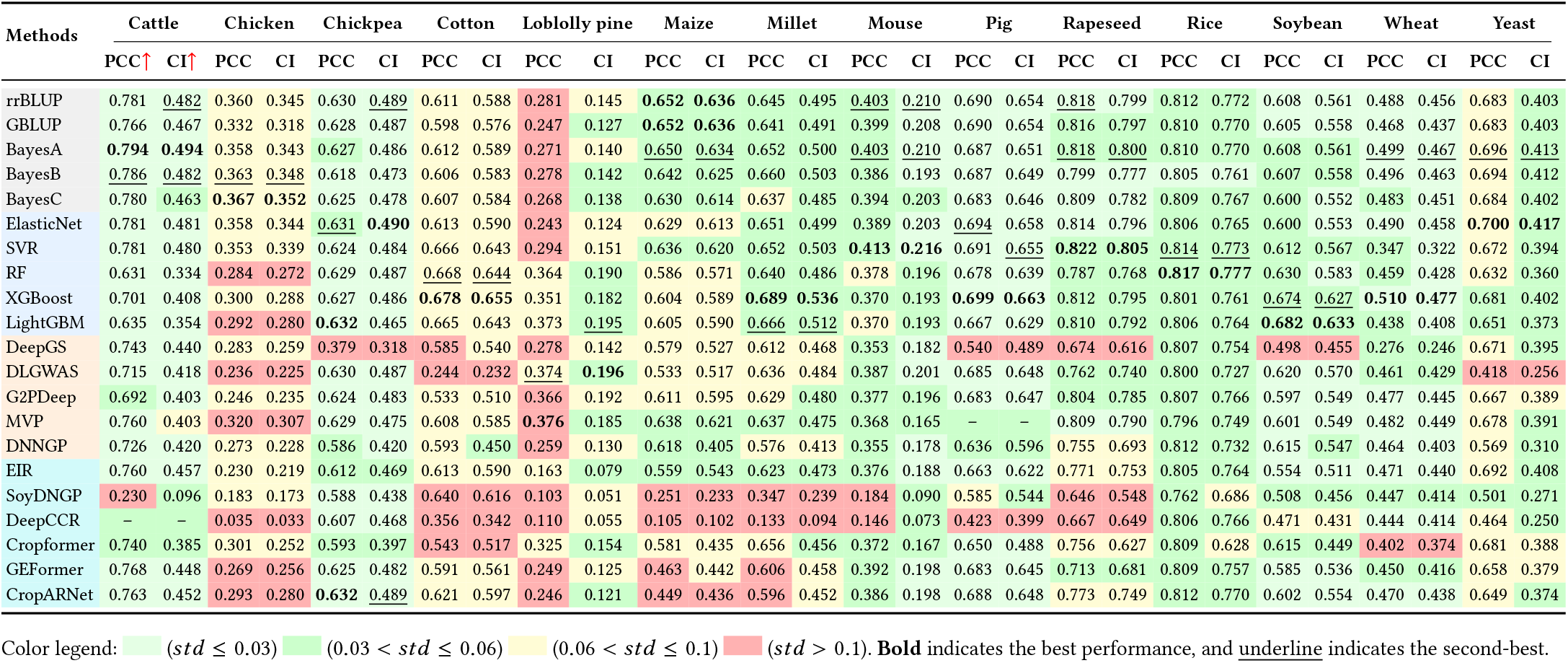
Pearson Correlation Coefficient (PCC) and Consistent Index (CI) of 21 Benchmark Methods on Various Species.

| Methods | Cattle |  | Chicken |  | Chickpea |  | Cotton |  | Loblolly pine |  | Maize |  | Millet |  | Mouse |  | Pig |  | Rapeseed |  | Rice |  | Soybean |  | Wheat |  | Yeast |  |
|---|---|---|---|---|---|---|---|---|---|---|---|---|---|---|---|---|---|---|---|---|---|---|---|---|---|---|---|---|
|  | PCC↑ | CI↑ | PCC | CI | PCC | CI | PCC | CI | PCC | CI | PCC | CI | PCC | CI | PCC | CI | PCC | CI | PCC | CI | PCC | CI | PCC | CI | PCC | CI | PCC | CI |
| rrBLUP | 0.781 | <u>0.482</u> | 0.360 | 0.345 | 0.630 | <u>0.489</u> | 0.611 | 0.588 | 0.281 | 0.145 | **0.652** | **0.636** | 0.645 | 0.495 | <u>0.403</u> | <u>0.210</u> | 0.690 | 0.654 | <u>0.818</u> | 0.799 | 0.812 | 0.772 | 0.608 | 0.561 | 0.488 | 0.456 | 0.683 | 0.403 |
| GBLUP | 0.766 | 0.467 | 0.332 | 0.318 | 0.628 | 0.487 | 0.598 | 0.576 | 0.247 | 0.127 | **0.652** | **0.636** | 0.641 | 0.491 | 0.399 | 0.208 | 0.690 | 0.654 | 0.816 | 0.797 | 0.810 | 0.770 | 0.605 | 0.558 | 0.468 | 0.437 | 0.683 | 0.403 |
| BayesA | **0.794** | **0.494** | 0.358 | 0.343 | 0.627 | 0.486 | 0.612 | 0.589 | 0.271 | 0.140 | <u>0.650</u> | <u>0.634</u> | 0.652 | 0.500 | <u>0.403</u> | <u>0.210</u> | 0.687 | 0.651 | <u>0.818</u> | <u>0.800</u> | 0.810 | 0.770 | 0.608 | 0.561 | <u>0.499</u> | <u>0.467</u> | <u>0.696</u> | <u>0.413</u> |
| BayesB | <u>0.786</u> | <u>0.482</u> | <u>0.363</u> | <u>0.348</u> | 0.618 | 0.473 | 0.606 | 0.583 | 0.278 | 0.142 | 0.642 | 0.625 | 0.660 | 0.503 | 0.386 | 0.193 | 0.687 | 0.649 | 0.799 | 0.777 | 0.805 | 0.761 | 0.607 | 0.558 | 0.496 | 0.463 | 0.694 | 0.412 |
| BayesC | 0.780 | 0.463 | **0.367** | **0.352** | 0.625 | 0.478 | 0.607 | 0.584 | 0.268 | 0.138 | 0.630 | 0.614 | 0.637 | 0.485 | 0.394 | 0.203 | 0.683 | 0.646 | 0.809 | 0.782 | 0.809 | 0.767 | 0.600 | 0.552 | 0.483 | 0.451 | 0.684 | 0.402 |
| ElasticNet | 0.781 | 0.481 | 0.358 | 0.344 | <u>0.631</u> | **0.490** | 0.613 | 0.590 | 0.243 | 0.124 | 0.629 | 0.613 | 0.651 | 0.499 | 0.389 | 0.203 | <u>0.694</u> | 0.658 | 0.814 | 0.796 | 0.806 | 0.765 | 0.600 | 0.553 | 0.490 | 0.458 | **0.700** | **0.417** |
| SVR | 0.781 | 0.480 | 0.353 | 0.339 | 0.624 | 0.484 | 0.666 | 0.643 | 0.294 | 0.151 | 0.636 | 0.620 | 0.652 | 0.503 | **0.413** | **0.216** | 0.691 | <u>0.655</u> | **0.822** | **0.805** | <u>0.814</u> | <u>0.773</u> | 0.612 | 0.567 | 0.347 | 0.322 | 0.672 | 0.394 |
| RF | 0.631 | 0.334 | 0.284 | 0.272 | 0.629 | 0.487 | <u>0.668</u> | <u>0.644</u> | 0.364 | 0.190 | 0.586 | 0.571 | 0.640 | 0.486 | 0.378 | 0.196 | 0.678 | 0.639 | 0.787 | 0.768 | **0.817** | **0.777** | 0.630 | 0.583 | 0.459 | 0.428 | 0.632 | 0.360 |
| XGBoost | 0.701 | 0.408 | 0.300 | 0.288 | 0.627 | 0.486 | **0.678** | **0.655** | 0.351 | 0.182 | 0.604 | 0.589 | **0.689** | **0.536** | 0.370 | 0.193 | **0.699** | **0.663** | 0.812 | 0.795 | 0.801 | 0.761 | <u>0.674</u> | <u>0.627</u> | **0.510** | **0.477** | 0.681 | 0.402 |
| LightGBM | 0.635 | 0.354 | 0.292 | 0.280 | **0.632** | 0.465 | 0.665 | 0.643 | 0.373 | <u>0.195</u> | 0.605 | 0.590 | <u>0.666</u> | <u>0.512</u> | 0.370 | 0.193 | 0.667 | 0.629 | 0.810 | 0.792 | 0.806 | 0.764 | **0.682** | **0.633** | 0.438 | 0.408 | 0.651 | 0.373 |
| DeepGS | 0.743 | 0.440 | 0.283 | 0.259 | 0.379 | 0.318 | 0.585 | 0.540 | 0.278 | 0.142 | 0.579 | 0.527 | 0.612 | 0.468 | 0.353 | 0.182 | 0.540 | 0.489 | 0.674 | 0.616 | 0.807 | 0.754 | 0.498 | 0.455 | 0.276 | 0.246 | 0.671 | 0.395 |
| DLGWAS | 0.715 | 0.418 | 0.236 | 0.225 | 0.630 | 0.487 | 0.244 | 0.232 | <u>0.374</u> | **0.196** | 0.533 | 0.517 | 0.636 | 0.484 | 0.387 | 0.201 | 0.685 | 0.648 | 0.762 | 0.740 | 0.800 | 0.727 | 0.620 | 0.570 | 0.461 | 0.429 | 0.418 | 0.256 |
| G2PDeep | 0.692 | 0.403 | 0.246 | 0.235 | 0.624 | 0.483 | 0.533 | 0.510 | 0.366 | 0.192 | 0.611 | 0.595 | 0.629 | 0.480 | 0.377 | 0.196 | 0.683 | 0.647 | 0.804 | 0.785 | 0.807 | 0.766 | 0.597 | 0.549 | 0.477 | 0.445 | 0.667 | 0.389 |
| MVP | 0.760 | 0.403 | 0.320 | 0.307 | 0.629 | 0.475 | 0.608 | 0.585 | **0.376** | 0.185 | 0.638 | 0.621 | 0.637 | 0.475 | 0.368 | 0.165 | – | – | 0.809 | 0.790 | 0.796 | 0.749 | 0.601 | 0.549 | 0.482 | 0.449 | 0.678 | 0.391 |
| DNNGP | 0.726 | 0.420 | 0.273 | 0.228 | 0.586 | 0.420 | 0.593 | 0.450 | 0.259 | 0.130 | 0.618 | 0.405 | 0.576 | 0.413 | 0.355 | 0.178 | 0.636 | 0.596 | 0.755 | 0.693 | 0.812 | 0.732 | 0.615 | 0.547 | 0.464 | 0.403 | 0.569 | 0.310 |
| EIR | 0.760 | 0.457 | 0.230 | 0.219 | 0.612 | 0.469 | 0.613 | 0.590 | 0.163 | 0.079 | 0.559 | 0.543 | 0.623 | 0.473 | 0.376 | 0.188 | 0.663 | 0.622 | 0.771 | 0.753 | 0.805 | 0.764 | 0.554 | 0.511 | 0.471 | 0.440 | 0.692 | 0.408 |
| SoyDNGP | 0.230 | 0.096 | 0.183 | 0.173 | 0.588 | 0.438 | 0.640 | 0.616 | 0.103 | 0.051 | 0.251 | 0.233 | 0.347 | 0.239 | 0.184 | 0.090 | 0.585 | 0.544 | 0.646 | 0.548 | 0.762 | 0.686 | 0.508 | 0.456 | 0.447 | 0.414 | 0.501 | 0.271 |
| DeepCCR | – | – | 0.035 | 0.033 | 0.607 | 0.468 | 0.356 | 0.342 | 0.110 | 0.055 | 0.105 | 0.102 | 0.133 | 0.094 | 0.146 | 0.073 | 0.423 | 0.399 | 0.667 | 0.649 | 0.806 | 0.766 | 0.471 | 0.431 | 0.444 | 0.414 | 0.464 | 0.250 |
| Cropformer | 0.740 | 0.385 | 0.301 | 0.252 | 0.593 | 0.397 | 0.543 | 0.517 | 0.325 | 0.154 | 0.581 | 0.435 | 0.656 | 0.456 | 0.372 | 0.167 | 0.650 | 0.488 | 0.756 | 0.627 | 0.809 | 0.628 | 0.615 | 0.449 | 0.402 | 0.374 | 0.681 | 0.388 |
| GEFormer | 0.768 | 0.448 | 0.269 | 0.256 | 0.625 | 0.482 | 0.591 | 0.561 | 0.249 | 0.125 | 0.463 | 0.442 | 0.606 | 0.458 | 0.392 | 0.198 | 0.683 | 0.645 | 0.713 | 0.681 | 0.809 | 0.757 | 0.585 | 0.536 | 0.450 | 0.416 | 0.658 | 0.379 |
| CropARNet | 0.763 | 0.452 | 0.293 | 0.280 | **0.632** | <u>0.489</u> | 0.621 | 0.597 | 0.246 | 0.121 | 0.449 | 0.436 | 0.596 | 0.452 | 0.386 | 0.198 | 0.688 | 0.648 | 0.773 | 0.749 | 0.812 | 0.770 | 0.602 | 0.554 | 0.470 | 0.438 | 0.649 | 0.374 |

Color legend: (*std* ≤ 0.03) (0.03 < *std* ≤ 0.06) (0.06 < *std* ≤ 0.1) (*std* > 0.1). **Bold** indicates the best performance, and <u>underline</u> indicates the second-best.

**Table 2.**
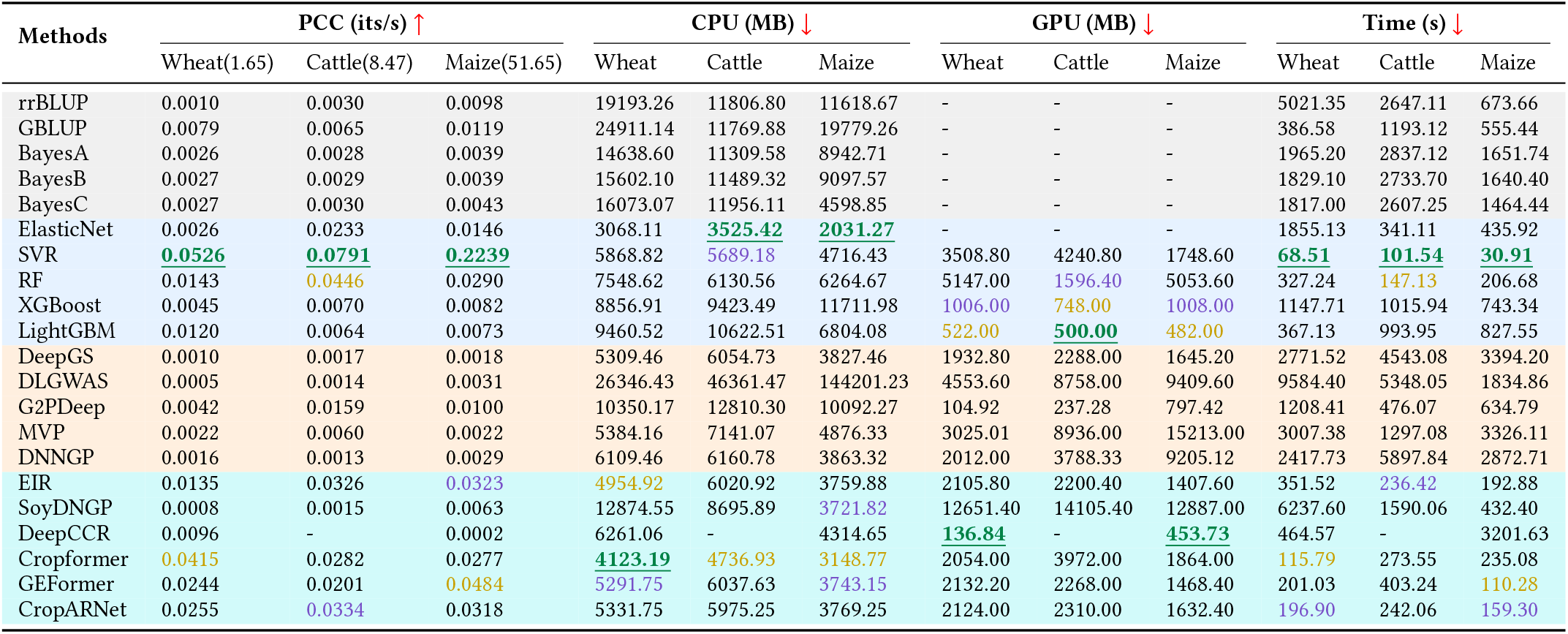
Model scalability comparison. The best, second-best, and third-best values are marked in green, orange, and purple.

As shown in Table 1, the experimental results show that statistical genetics–based methods exhibit stable and competitive predictive performance across most species, indicating that additive genetic effects remain the primary effective source of information in genomic prediction tasks. This also supports the rationality and reliability of using statistical genetic methods such as rrBLUP and Bayesian approaches as baselines in existing studies. Among machine learning methods, tree-based models (e.g., XGBoost and Light-GBM) outperform statistical genetic approaches in some species, demonstrating their ability to capture limited nonlinear relationships. CNN-based methods show larger performance variability across species: while high accuracy can be achieved in certain tasks (e.g., DeepGS), their overall stability is strongly influenced by data scale and assumptions about local genomic modeling. Hybrid-architecture methods do not achieve the best overall performance; however, they approach or even outperform statistical methods on multiple species. Their performance across species is characterized by a ‘high ceiling with high variance’ pattern—exhibiting strong representational capacity in some scenarios, but with clearly insufficient stability and consistency in generalization.

#### 4.1.2 Analysis of Genetic Structure Differences

In the Cattle dataset (*P*/*N* = 8.47), the three traits exhibit distinct genetic architectures: fat percentage (fpro) is driven by one major gene with many small-effect loci; (2) milk yield (mkg) by a few moderate-effect loci plus numerous small effects; (3) somatic cell score (scs) primarily by highly dispersed small-effect loci (see Appendix B, Fig. S3). Benchmarking 21 methods reveals a consistent high–medium–low performance ranking across the three traits, suggesting that model architecture is not the primary determinant of predictive performance. Motivated by this observation, we conduct feature ablation experiments by removing the major-effect features of fpro, medium–small effects of mkg, and partial small effects of scs, to investigate the underlying cause of this global pattern.

Figure 4 shows that performance differences across traits arise from variations in the dimensionality and compressibility of predictable genetic information. For fpro, the information is highly concentrated in a low-dimensional main-effect subspace, and relying solely on the main-effect features can nearly recover the original performance, while removing this set leads to a substantial drop. In contrast, removing or using only the corresponding features in mkg and scs has a limited impact, indicating that overlapping loci across traits do not necessarily correspond to transferable predictive information. The mkg exhibits a dual-source structure of ‘few highly identifiable effects + many background weak effects’, while scs displays highly dispersed and redundant polygenic features. These results suggest that models can optimize how to aggregate available information, but cannot surpass the limits imposed by the inherent structure of genetic information.

**Figure 4.**
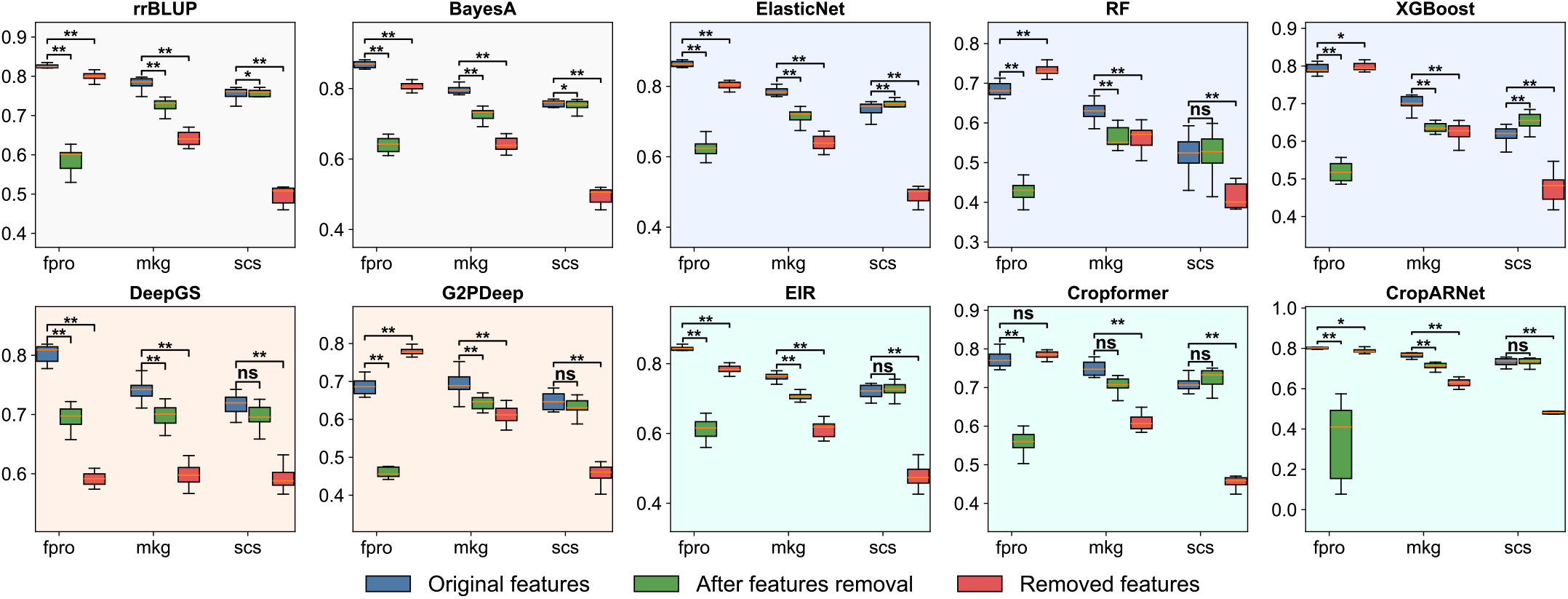
Performance of representative methods across traits with different genetic architectures. Original features use all features; after features removal excludes frpo identified major-effect features; removed features use only removal features. Statistical significance: \*\**p <* 0.01, \**p <* 0.05, *ns*, not significant.

### 4.2 Sensitivity analysis

To evaluate the sensitivity of GP, we conducted a systematic sub-sampling study to analyze the performance variations of different GP algorithms under varying sample sizes, feature dimensions, and random initialization conditions. Specifically, for the wheat dataset on grain yield (*P*/*N* = 1.65), we considered five training sample sizes (1,000, 2,000, 4,000, 6,000, and 8,000). For each sample size, three different random seeds (42, 123, and 246) were used for data splitting to assess model scalability and robustness under different sampling conditions. Model performance was primarily evaluated using PCC and Normalized Mean Absolute Error (NMAE). In addition, we analyzed GP model performance on the mouse body weight trait (*P*/*N* = 5.77) under varying feature dimensions (300, 1,000, 3,000, 5,000, and 10,000) and different random initialization conditions (42, 123, and 246) (see Appendix B.2, Fig. S6).

As shown in Figure 5, overall, most GP algorithms exhibit a steady performance improvement as the sample size increases, with smooth and consistent trends across different sampling scales. This observation aligns with expectations and indicates that increasing the number of training samples effectively enhances predictive accuracy and generalization ability. From the perspective of sensitivity to random seeds, most models are influenced by data partitioning under small-sample conditions. In contrast, XGBoost and Light-GBM demonstrate high consistency with relatively low variance, suggesting strong stability with respect to data subsampling in small-sample regimes. By comparison, several deep learning models (e.g., DeepGS, DLGWAS, SoyDNGP, and Cropformer) exhibit higher sensitivity to random initialization when training data are limited, as evidenced by pronounced fluctuations in PCC and NMAE curves and larger error ranges. This indicates a stronger dependence on training sample size and randomness. Notably, other deep learning models, such as EIR, GEFormer, and CropARNet, show improved stability and consistency across different sample sizes. Their performance not only improves substantially with increasingsample size but also becomes less sensitive to random seed variations, reflecting stronger data modeling capacity and predictive performance. These results suggest that more effective model designs can partially alleviate the instability of deep learning models in small-sample GP scenarios.

**Figure 5.**
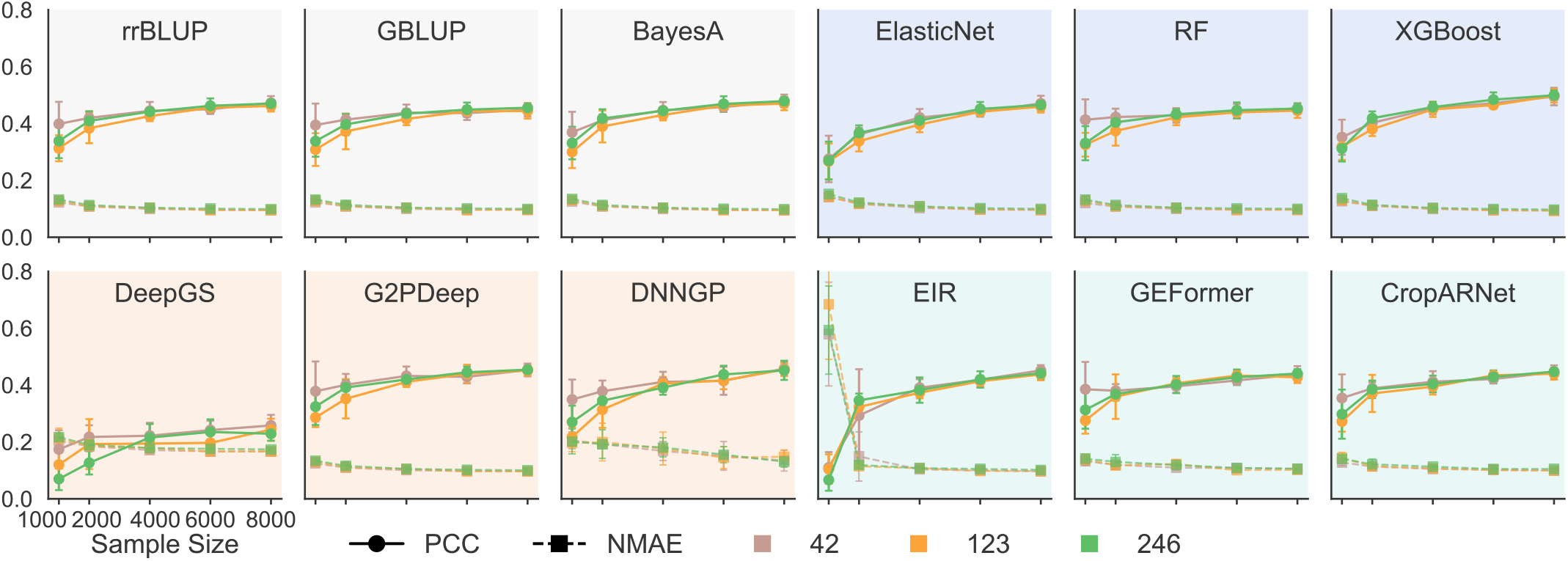
Performance of GP models in terms of PCC and NMAE (Normalized Mean Absolute Error) under different sample sizes and random seed splits. Circles represent PCC values, squares denote NMAE values, and different colors correspond to different random seeds.

### 4.3 Robustness analysis

To evaluate the robustness of GP algorithms under scenarios that more closely resemble real-world biological sequencing errors, we introduced random noise at the genotype level for the Yeast magnesium_sulfate trait (*P*/*N* = 11.53, normal distributions) and the Rice grain weight trait (*P*/*N* = 2.53, non-normal distributions) by randomly flipping a certain proportion of genotype loci. The noise proportions were set to 0.05, 0.1, 0.2, 0.3, and 0.5 (Fig. 6). In addition, we evaluated the impact of missing on the Pig back fat thickness trait (*P*/*N* = 7.82, normal distributions) and the Rapeseed Flower Time trait (*P*/*N* = 10.03, non-normal distributions), with missing intensities set to 0.1, 0.3, and 0.5 (Fig. S8).

**Figure 6.**
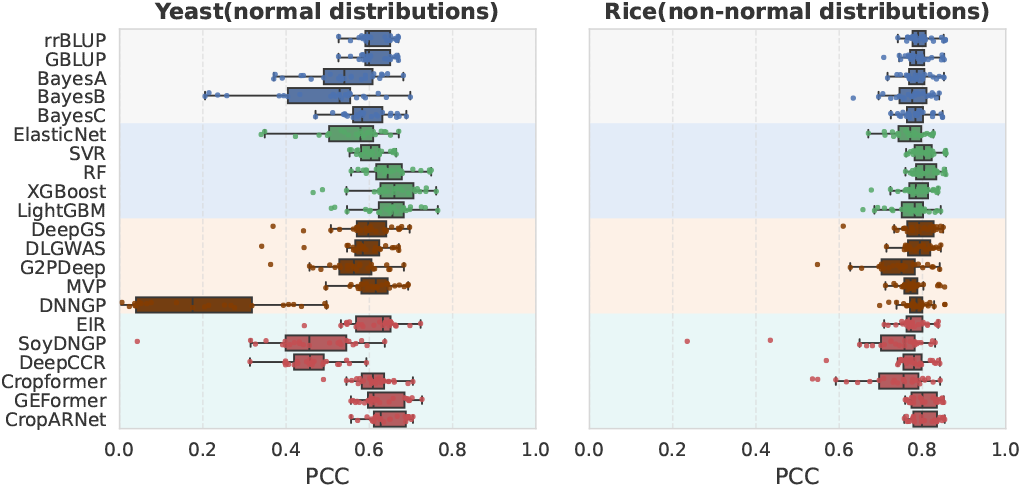
Boxplot of PCC values for 21 GP algorithms under different noise levels. A larger spread of the boxplot indicates a bigger performance variability under noise perturbations.

Overall, as the noise level increases, the predictive performance (PCC) of most methods exhibits a general downward trend, accompanied by increased variance (*R*^2^ result see Appendix B.3, Fig.S7). This trend is particularly pronounced in the Yeast dataset. Statistical genetics methods and several machine learning approaches (e.g., rrBLUP, GBLUP, SVR, and RF) demonstrate relatively stable performance, indicating a low sensitivity to random single-locus errors. This robustness can be attributed to their reliance on genome-wide aggregated information and the regularization effects inherent in these models. In contrast, GP methods based on convolutional and hybrid architectures exhibit substantial within-group performance divergence. In particular, CNN-based approaches heavily depend on the extraction of local sequence patterns and are therefore more vulnerable to the accumulation of locus-level errors, which can disrupt local pattern learning. Nevertheless, across both distributional settings, GEFormer and CropARNet consistently show the most compact PCC distributions across varying noise levels, outperforming statistical genetics and traditional machine learning methods and demonstrating relatively strong robustness

### 4.4 Scalability analysis

To systematically evaluate the scalability and computational efficiency of different GP models in practical applications, we conducted experiments on three representative benchmark datasets: wheat (*P*/*N* = 1.65), cattle (*P*/*N* = 8.47), and maize (*P*/*N* = 51.65). We assessed model performance from multiple perspectives, including prediction performance achieved per unit time (measured as PCC per second), CPU and GPU memory consumption, and total runtime, with the aim of comprehensively characterizing the tradeoffs between predictive efficiency and computational cost across various GP models. The experimental platform sees Appendix A.4. The experimental results reveal substantial differences in computational efficiency and scalability among different categories of GP models. SVR achieves relatively high predictive gains per unit time across all three datasets while maintaining manageable computational costs, demonstrating a favorable balance between efficiency and performance. Cropformer, CropARNet, and GEFormer also exhibit low memory consumption and stable runtimes across the three datasets; although their predictive gains per unit time are comparatively limited, their overall computational overhead remains relatively small. Notably, traditional statistical models do not perform competitively in this evaluation, often incurring higher CPU memory consumption and longer runtimes, with pronounced efficiency variability observed among different methods. In summary, genetic statistical models have limited scalability under resource-constrained or extremely large-scale data scenarios, whereas DL and hybrid ML models can balance efficiency and performance and exhibit greater scalability.

### 4.5 Online Application Platform

We further developed an online application platform for GPBench, which comprises three modules: intelligent data analysis (Agent), benchmark result visualization (Analysis), and user task testing (Tasks), providing services to the research community. For the Agent module, we conducted case-based usage analyses, including a result interpretation task (see Appendix B.4, Fig. S9) and an algorithm recommendation task (Figure 7). When users pose the request “Could you recommend the most suitable algorithm for this dataset?”, the Agent interprets user requirements, searches for the most similar datasets in the benchmark experience (e.g., Wheat, Pig), and then identifies the most suitable algorithm XGBoost, and provides the reason. The agent reports, “The upstream answer explicitly recommends XGBoost as the most suitable algorithm for the dataset based on its superior performance in accuracy, robustness, computational efficiency, memory usage, and generalizability.”

**Figure 7.**
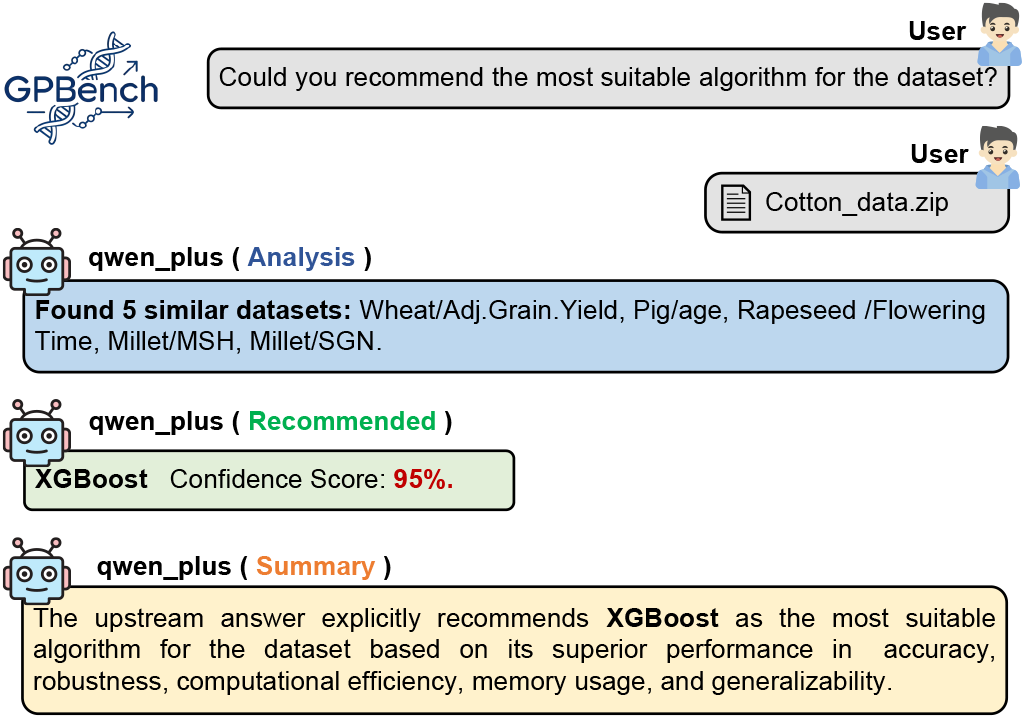
User and agent chat process in the algorithm recommendation task.

## 5 Discussion

To bridge the gap between the rapid development of GP methods and the lack of systematic evaluation, we propose GPBench, a systematic and scalable benchmarking framework for genomic prediction. We evaluate 21 GP methods from the perspectives of effectiveness, sensitivity, robustness, and scalability. Experimental results show that statistical genetics–based methods exhibit stable and competitive predictive performance across most species. Feature engineering–based machine learning methods, especially tree-based models, outperform statistical genetic methods on certain species, demonstrating their ability to capture limited nonlinear relationships. CNN-based methods and hybrid architecture–based methods also achieve performance comparable to or even exceeding that of statistical genetic methods on multiple species; however, their stability and generalization consistency remain clearly insufficient. At present, GPBench does not yet include prediction methods that incorporate environmental or multi-omics information. In future work, we plan to extend the GPBench platform by further integrating multimodal datasets (such as transcriptomics, epigenomics, and environmental data) and by updating state-of-the-art methods. Meanwhile, we sincerely welcome feedback and suggestions from the research community to jointly promote the development of GP methods.

## 6 Conclusion

We propose GPBench, a systematic and scalable benchmark framework for genomic prediction. GPBench provides a unified evaluation of 21 state-of-the-art methods across datasets covering 16 species and 57 trait scenarios, from four complementary dimensions. In addition, we leverage an agent analysis to interpret and synthesize the benchmarking results. Finally, we release the GPBench benchmarking platform and GPBench library, expect GPBench to promote their broader application and advancement in life science research.

## A Appendix

### A.1 Supplementary Benchmark Dataset

#### A.1.1 Details of Benchmark Dataset

1. **Cattle** [58]: This dataset consists of 5,024 German Holstein cattle. After genotyping and quality control, 42,551 SNPs were retained for subsequent genomic prediction. Phenotypic records include three traits: milk fat percentage, milk yield, and somatic cell score.
2. **Chicken** [8]: This dataset comprises 1,063 chicken samples. The original genotype data were subjected to quality control by removing SNPs with minor allele frequency (MAF) < 0.01 or genotype missing rate > 0.01, followed by linkage disequilibrium (LD) pruning, resulting in 4,293 SNPs retained. For phenotypes, egg weight data at two time points with the lowest missing rates, 28 weeks of egg weight and 36 weeks of egg weight, were used.
3. **Chickpea** [6, 45]: This dataset includes 2,921 chickpea accessions. After quality control of the original genotypes, SNPs with MAF < 0.05 or genotype missing rate > 0.01 were removed, followed by LD pruning, yielding a total of 53,193 SNPs. Three phenotypic traits with the lowest missing rates were retained: days to 50% flowering, plant height, and hundred-seed weight.
4. **Cotton** [6, 16]: This dataset contains 1,245 cotton accessions. After genotype quality control, SNPs with MAF < 0.05 or genotype missing rate > 0.01 were removed, leaving 162 SNPs for analysis. Phenotypic traits used in this study include fiber length, fiber strength, fiber micronaire, and fiber elongation rate.
5. **Horse** [58]: This dataset consists of 480 horses samples. After genotyping and quality control, 48,697 SNPs were retained for genomic prediction. Phenotypic data were adopted directly from the published study, where coat color traits were encoded as 1, 2, 3, and 4, representing chestnut, gray, dun, and black/bay, respectively.
6. **Human** [49]: The dataset includes two diseases: breast cancer and age-related macular degeneration. The breast cancer dataset contains 4,938 samples with 5,607 SNPs, while the age-related macular degeneration dataset contains 146 samples with 103,611 SNPs. The phenotype data are discrete, where 0 indicates non-disease and 1 indicates disease.
7. **Loblolly Pine** [39]: This dataset consists of 926 samples. We used the genotype dataset processed and released in the corresponding publication, 4,853 SNPs were retained for genomic prediction. For the 17 traits, phenotypes with high missing rates were filtered out, and data for 7 traits were retained for subsequent analysis. Phenotypic traits used in this study include root number, presence or absence of roots, 5-and 6-carbon sugar content, wood specific gravity, latewood percentage at year 4, lignin content, and tree stiffness.
8. **Maize** [6, 53]: This dataset consists of 1,428 maize samples derived from maternal inbred lines. The original genotype data were subjected to quality control by removing SNPs with MAF < 0.05 or genotype missing rate > 0.01, followed by LD pruning, resulting in 75,304 SNPs retained. Phenotypic traits used in this study include days to tasseling, plant height, and ear weight.
9. **Millet** [6, 50]: This dataset includes 827 foxtail millet accessions. After quality control of the original genotypes by removing SNPs with MAF < 0.05 or genotype missing rate > 0.01, a total of 20,967 SNPs were retained. Phenotypic traits include six growth traits: top second leaf length, top second leaf width, main stem height, main stem width, panicle diameter of the main stem, and fringe neck length; and six yield-related traits: panicle length of the main stem, per plant grain weight, main stem panicle weight, hundred kernel weight, spikelet number of the main stem, and grain number per spike.
10. **Mouse** [44]: For the model organism mouse dataset, 1,762 outbred mice were retained after filtering, and 10,168 SNP genotypes curated by the original publication were used [44] Phenotypic traits include body weight, anxiety-like behavior, and body mass index.
11. **Pig** [55]: This dataset consists of 4,157 pig samples. The original genotype data were subjected to quality control by removing SNPs with MAF < 0.05 or genotype missing rate > 0.01, followed by LD pruning, resulting in 32,508 SNPs retained. Two phenotypic traits were used: age at 100 kg body weight and backfat thickness at 100 kg body weight.
12. **Rapeseed** [6, 52]: This dataset consists of 991 rapeseed samples. After genotype quality control by removing SNPs with MAF < 0.05 or genotype missing rate > 0.01, 9,944 SNPs were retained. Flowering time was used as the phenotypic trait.
13. **Rice** [6, 21]: This dataset includes 705 rice accessions. After quality control of the original genotypes by removing SNPs with MAF < 0.05 or genotype missing rate > 0.01, 1,785 SNPs were retained. Phenotypic traits include grain length, grain weight, 1000-Grain weight, heading date, plant height, number of productive panicle per plant, and grain yield per plant.
14. **Soybean** [6, 24]: This dataset consists of 2,795 soybean accessions. After genotype quality control by removing SNPs with MAF < 0.05 or genotype missing rate > 0.01, followed by LD pruning, 51,442 SNPs were retained. Phenotypic traits include beginning bloom date, plant height, and valid branch number.
15. **Wheat** [34]: This dataset contains 10,375 wheat samples. We directly used 17,113 SNPs curated by the original publication [34]. Phenotypic trait is adjusted grain yield.
16. **Yeast** [41]: This dataset includes 1,008 yeast samples and 11,623 SNPs curated by the original publication [41]. Phenotypic traits with the lowest missing rates were selected, including growth responses under cobalt chloride, magnesium sulfate, and tunicamycin treatments.

#### A.1.2 Phenotype Kernel Density Estimation

For benchmark datasets, kernel density estimation was performed on the phenotypes to explore the distribution patterns across different species, and the results are shown in Figure S1. We observed that the traits of most species exhibit unimodal distributions that are approximately normal, which is consistent with the genetic characteristics of quantitative traits, namely that they are controlled by multiple genes and display continuous variation. In addition, bimodal distributions were observed for certain traits in some species, such as grain length in rice and flowering time in rapeseed. This suggests the presence of population structure differences in the collected datasets, where distinct genetic backgrounds or environmental factors influence the phenotype distributions, and it also reflects the inherent complexity of genomic prediction tasks.

### A.2 Details of Benchmark Algorithms

**rrBLUP** [10]: is a genomic prediction method based on the linear mixed model. It assumes that all SNP effects follow the same normal distribution and applies uniform shrinkage to the effects through ridge regression, thereby preventing overfitting. This method is one of the most classical and widely used baseline models in genomic selection.

**GBLUP** [13]: is also a linear genomic prediction model. Similar to rrBLUP, GBLUP assigns the same variance to all loci and characterizes the genetic similarity among individuals through the genomic relationship matrix.

**BayesA** [31]: is a Bayesian genomic prediction model. Unlike rrBLUP, it assumes that each SNP effect follows a *t*-distribution with an independent variance. This formulation allows SNPs to have effects of different magnitudes and provides a more flexible description of SNP effect heterogeneity compared with rrBLUP.

**BayesB** [31]: is an extension of BayesA. In practice, the distribution of genetic variance across loci is such that many loci contribute no genetic variance, whereas only a small proportion have non-zero effects. To better reflect this reality, BayesB introduces a variable selection mechanism by assuming that only a subset of SNPs contribute to the trait, while the remaining SNP effects are set to zero.

**BayesC**[14]: is a further extension of BayesB, in which all nonzero SNP effects are assumed to share a common variance, thereby reducing model complexity. BayesC retains the variable selection capability while offering improved computational efficiency.

**ElasticNet** [62]: was proposed for the analysis of high-dimensional data by combining the *L*_1_ (Lasso) and *L*_2_ (ridge regression) penalty terms. The *L*_1_ component enables automatic variable selection among SNPs, while the *L*_2_ component promotes grouped selection and stabilizes the solution path under random sampling, thereby improving the accuracy of genomic prediction.

**SVR** [28]: is a classical machine learning method that models genetic effects by mapping genotype data into a high-dimensional feature space through the use of kernel functions. SVR does not rely on explicit genetic assumptions and shows advantages in handling complex and non-additive genetic architectures; however, it is sensitive to parameter selection, such as the choice of kernel function and penalty coefficient.

**RF** [4]: is an ensemble learning method based on multiple decision trees. By employing bootstrap sampling and random feature selection, RF effectively reduces model variance. It is capable of automatically capturing nonlinear relationships and high-order interaction effects and exhibits strong robustness to noise and outliers.

**XGBoost** [7]: is a machine learning method based on gradient boosting decision trees, which improves predictive performance by iteratively fitting residuals. It performs well in modeling complex nonlinear patterns and effectively controls overfitting through regularization mechanisms, demonstrating strong generalization ability in high-dimensional feature spaces and large-scale datasets.

**LightGBM** [54]: is a machine learning framework developed by Microsoft for large-scale data analysis. It adopts histogram-based splitting and a leaf-wise tree growth strategy, which significantly improves training efficiency. While maintaining high predictive accuracy, LightGBM reduces computational resource consumption and is particularly suitable for genomic prediction tasks involving large sample sizes and high-dimensional marker data.

**Figure S1.**
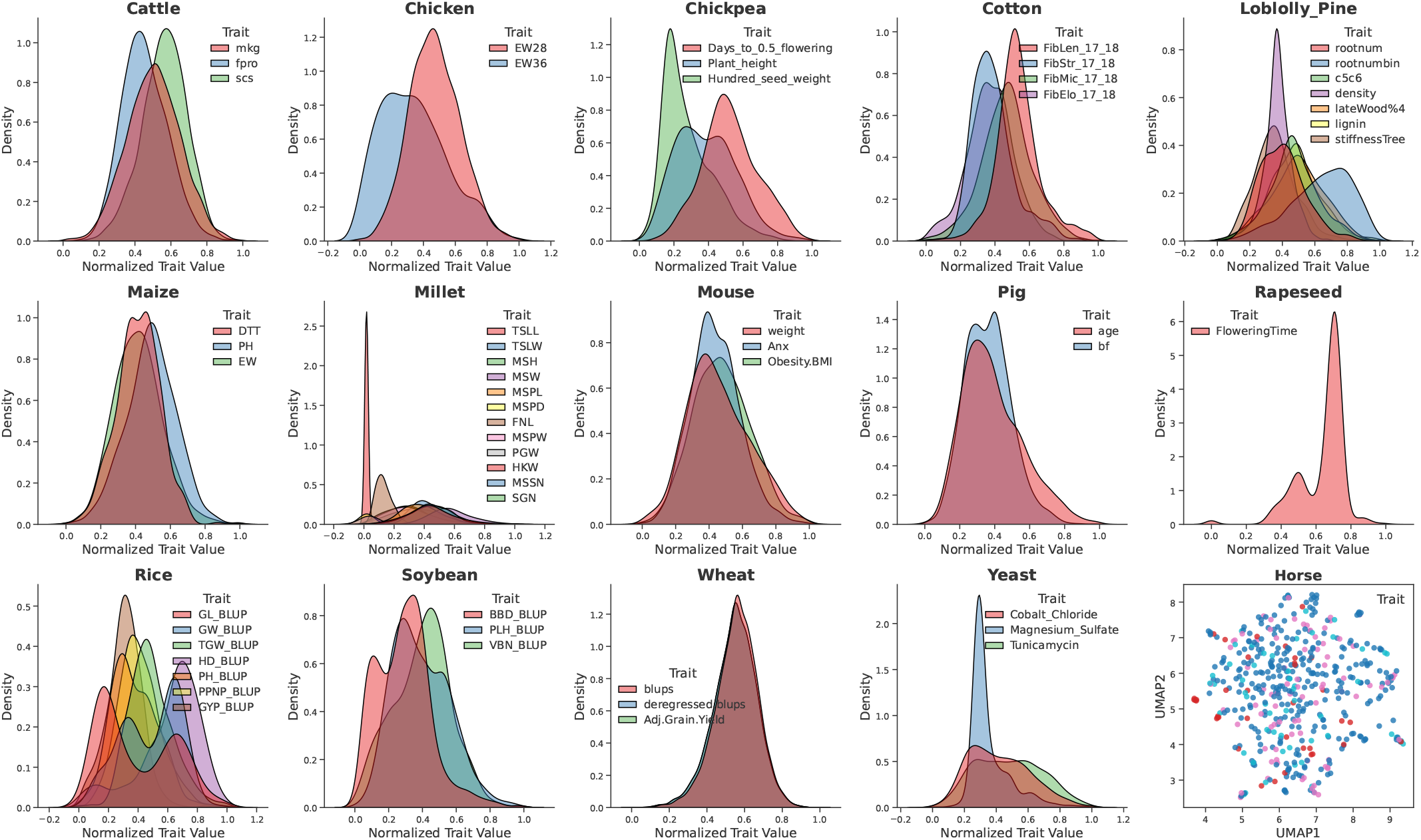
Kernel density estimation plots of continuous phenotypes and scatter plots of discrete phenotypes across multiple species datasets.

**DeepGS** [29]: is a deep learning method based on convolutional neural networks(CNN), consisting of an input layer, convolutional layers, pooling layers, fully connected layers, and an output layer. During prediction, latent variables are used to jointly represent genotypic features, thereby reducing the complexity of high-dimensional genotype data. To prevent overfitting, an early stopping mechanism is incorporated during the training process.

**DLGWAS** [26]: is a deep learning method based on a dual-stream convolutional neural network. SNPs are first fed into two parallel convolutional streams for feature extraction, after which their outputs are aggregated through a summation layer and subsequently passed to a single convolutional layer and then to fully connected layers for accurate phenotype prediction. In addition, dropout layers and an early stopping mechanism are employed to ensure effective model optimization.

**G2PDeep** [59]: is also a deep learning method based on a dual-stream CNN. The model consists of a dual-CNN module followed by a fully connected neural network. After one-hot encoding, the input is fed into the dual CNN, and then passed through a single CNN to further enhance marker representations. The integrated marker representations are flattened and fed into an output layer to estimate quantitative phenotypes.

**MVP** [36]: is a method based on deep residual neural networks. The model consists of two residual blocks composed of convolutional filters and activation layers, followed by two fully connected layers with sigmoid outputs. The convolutional layers in ResNet can extract hierarchical features and nonlinear local patterns from image or sequence data, making the model well-suited for large-scale genomic prediction tasks.

**DNNGP** [48]: is also a CNN-based deep learning model that stacks multiple linear and nonlinear processing units in a hierarchical manner. Specifically, DNNGP comprises three CNN layers for feature extraction, one batch normalization layer to address internal covariate shift, and two dropout layers together with an early stopping strategy to prevent overfitting. Furthermore, to accelerate learning, improve convergence speed, and alleviate the vanishing gradient problem, DNNGP employs the rectified linear unit (ReLU) activation function.

**EIR** [40]: is a deep learning framework for polygenic risk scoring based on gated linear networks (GLNs). Specifically, the GLN feature extractor replaces convolutional layers with locally connected layers (LCLs) to capture position-specific and non-additive genetic effects. The LCLs use group-wise sparse connections, which substantially reduce the number of parameters compared with fully connected layers.

**SoyDNGP** [11]: is a CNN-based deep learning method. Unlike the classical three-layer convolutional architecture, SoyDNGP adopts the block segmentation concept from visual geometry group networks, in which each convolutional block consists of a convolutional layer, a normalization layer, and a ReLU activation layer. In addition, coordinate attention (CA) modules are introduced after the first and the last convolutional layers. This strategy enhances the modeling of positional information within feature matrices and across channels, thereby improving the extraction of spatial information.

**DeepCCR** [30]: is a deep learning method that combines convolutional neural networks with bidirectional long short-term memory (BiLSTM) networks. Specifically, DeepCCR first performs feature extraction using one-dimensional convolutional layers, followed by a max-pooling layer, a BiLSTM layer, another one-dimensional convolutional layer, a flattening layer, and two fully connected layers. Finally, a fixed-dimensional feature vector is produced by the output layer to generate the final prediction.

**Cropformer** [47]: is a deep neural network that integrates convolutional neural networks with multi-head self-attention mechanisms. In this framework, convolutional layers in the CNN automatically extract features from raw input data and transform them into informative representations. These features are then fed into an attention module to generate decision vectors for prediction.

**GEFormer** [56]: is a genomic prediction method designed to model genotype–environment interactions. For ease of comparison, only the genomic feature extraction module of GEFormer is used in this study for phenotype prediction. This module employs gMLP to extract SNP features. gMLP is a simple network architecture based on multilayer perceptrons (MLPs) with a gating mechanism, which enables feature extraction at both local and global levels, thereby capturing long-range dependencies among SNPs.

**CropARNet** [61]: is a deep learning method that integrates self-attention mechanisms with deep residual networks. Specifically, the model architecture consists of an attention module and a residual network. First, the attention module uses multi-layer fully connected networks to generate attention weights for each SNP. These weights are then fed into the residual network and element-wise multiplied with the original genomic features to obtain enhanced representations. Finally, the refined representations are passed to a regression layer to produce the predicted phenotypic values.

### A.3 Evaluation Metrics

To comprehensively evaluate the performance of GP models, this study employs the Pearson correlation coefficient (PCC), mean absolute error (MAE), coefficient of determination (*R*^2^), root mean squared error (RMSE), and consistency index (CI) [51] as evaluation metrics for regression tasks; the Accuracy, Precision, Recall, and F1-score as evaluation metrics for classification tasks.

#### Pearson Correlation Coefficient (PCC)

The PCC is used to measure the linear correlation between the predicted and the groundtruth phenotypic values. Its value ranges from 1 to 1, where a larger PCC indicates a stronger consistency between predictions and truths. The PCC is calculated as follows:

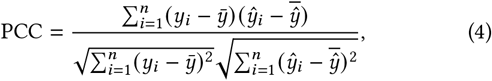

where *y*_*i*_ denotes the ground-truth phenotypic value of the *i*-th sample, *ŷ*_*i*_ represents the corresponding predicted value, 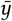 and 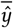 are the mean values of the ground-truth and predicted phenotypes, respectively, and *n* is the number of samples.

#### Mean Absolute Error (MAE)

The MAE measures the average magnitude of the errors between the predicted and the ground-truth values, reflecting the numerical prediction accuracy. It is defined as:

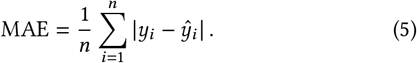

##### Coefficient of Determination (*R*^2^)

The *R*^2^ evaluates the proportion of phenotypic variance explained by the GP model. A larger *R*^2^ indicates better explanatory power of the model. It is defined as:

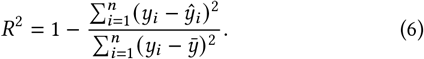

where 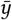 denotes the mean of the ground-truth phenotypic values.

#### Root Mean Squared Error (RMSE)

The RMSE is obtained by squaring the prediction errors, averaging them, and then taking the square root, which penalizes larger errors more heavily and reflects the overall prediction stability of the model. It is calculated as:

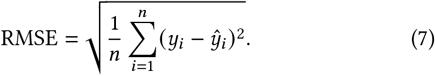

#### Consistency Index (CI)

The PCC measures the consistency between predicted and true phenotypic values in terms of trends, while MAE reflects the magnitude of numerical prediction errors. However, using PCC or MAE alone is insufficient to comprehensively evaluate the overall predictive performance of a model. Therefore, we introduce a composite metric CI, which jointly characterizes trend consistency and numerical accuracy [51]. The CI is defined as:

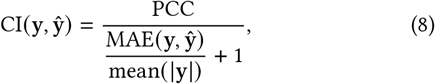

where PCC denotes the Pearson correlation coefficient between the predicted and ground-truth phenotypic values, MAE (y, ŷ) represents the mean absolute error on the test set, and mean (|y|) is the mean of the absolute values of the true phenotypes in the test set. In practice, CI values are typically in 0, 1, with larger values indicating better overall predictive performance. When the CI approaches 1, the predicted results exhibit high trend consistency (PCC close to 1) and small numerical errors (MAE close to 0). The inclusion of mean (|y|) as a normalization coefficient ensures that the CI remains within a reasonable range and enhances comparability across different traits or datasets. Consequently, CI serves as an effective indicator for GP model evaluation and selection.

#### Accuracy

The Accuracy is used to measure the overall correctness of GP model predictions in classification tasks, defined as the proportion of individuals whose predicted phenotypes match their true phenotypes. Its calculation formula is:

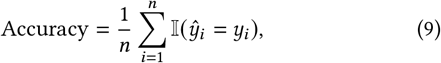

where *I*(·) is the indicator function, equal to 1 if the prediction is correct and 0 otherwise.

#### Precision

The Precision is used to measure the proportion of individuals predicted as a specific target phenotype (positive class) that actually possess that phenotype. It is calculated using macro-averaging, with the formula:

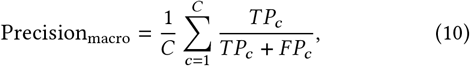

where *C* denotes the number of phenotype classes, and *TP*_*c*_ and *FP*_*c*_ represent the numbers of true positives and false positives for class *c*, respectively.

#### Recall

The Recall is used to measure the proportion of individuals with the target phenotype that are successfully predicted by the model. It is also calculated using macro-averaging, with the formula:

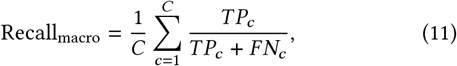

where *F N*_*c*_ denotes the number of false negatives for class *c*.

#### F1-score

The F1-score is the harmonic mean of Precision and Recall, especially suitable for genetic data with imbalanced phenotype distributions. Its calculation formula is:

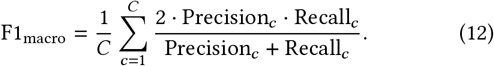

### A.4 Experimental Settings

#### A.4.1 Experimental Environment

All experiments were conducted on a Linux-based server equipped with a 128-core Intel CPU, 197 GB of system memory, and a single NVIDIA L40 GPU with 45 GB of GPU memory. The CUDA version used was 12.2. The software environment was built using Conda, with Python version 3.9.23. All deep learning models were implemented using PyTorch 2.9.1, together with torchvision 0.24.1. In addition, auxiliary libraries such as Optuna 4.6.0 were employed. All experiments were executed under the same hardware and software configurations to ensure the reproducibility and fairness of the experimental results.

#### A.4.2 Evaluation Protocol

To ensure a fair comparison of different algorithms under a unified benchmark, we adopt a standardized implementation and evaluation protocol for all 21 GP methods. Our overall objective is to **standardize and refactor the open-source implementations without altering the core modeling assumptions or network architectures of the original methods**, such that they satisfy the following requirements: (i) unified data interfaces; (ii) unified cross-validation strategy and evaluation metrics; (iii) unified output formats; and (iv) experimental reproducibility.

i. *Unified data interface:* Each species (or dataset) is organized into a dedicated directory. Each dataset mainly consists of the following files. **genotype**: denoted as *X* ∈ ℝ^*n*×*p*^, where *n* is the number of samples and *p* is the number of SNPs; **phenotype**: denoted as *Y* ∈ ℝ^*n*×*t*^, where *t* is the number of phenotypes. For any phenotype *j* the corresponding label vector is defined as *y* = *Y* [:, *j*]. Missing values are imputed using the mean value computed from the training set.
ii. *Unified cross-validation and evaluation metrics:* To ensure fairness and robustness in performance evaluation, we adopt a 10-fold cross-validation strategy for all datasets. Before splitting, each dataset is randomly shuffled using a fixed random seed (*randomseed* = 42). All compared methods share identical data partitions to guarantee a fair comparison. For models involving hyperparameters, Optuna is used for automated hyperparameter optimization. Specifically, we employ a nested cross-validation scheme: in the inner loop, Optuna performs cross-validation and selects the optimal hyperparameters based on the Pearson correlation coefficient (PCC) or the F1-score; the outer 10-fold cross-validation is then used to obtain unbiased performance estimates.
iii. *Unified output format:* To facilitate subsequent statistical analysis, we standardize the output of experimental results for each method–species–phenotype combination as follows. Per-fold predictions containing two columns (*Y* _*test* and *Y* _*pred*); Console or log summaries reporting the 10-fold average PCC, MAE, MSE, and *R*^2^, along with the runtime and the peak CPU/GPU memory consumption.
iv. *Experimental reproducibility:* To ensure experimental reproducibility, we fix the random seed for all methods. For PyTorchbased implementations, deterministic computation settings are enabled. For implementations based on *scikit* ™ *learn, cuML*, or R, the corresponding *random*_*state* or equivalent parameters are also fixed.

### A.5 Hyperparameter Settings

The GPBench framework currently integrates 21 benchmark methods involving different numbers of hyperparameters. Among them, genetic statistical methods do not require predefined hyperparameters, whereas feature engineering–based machine learning methods, convolution-based deep learning models, and hybrid architecturebased deep learning models require the specification of a large number of hyperparameters. To ensure a fair comparison of their true performance in genomic prediction tasks, and considering the broad range of datasets evaluated, the original literature does not cover all datasets used in this study. Therefore, we adopted the following strategy for hyperparameter configuration. For methods requiring hyperparameter optimization, we employ a nested cross-validation scheme, where the outer loop is used for an unbiased estimation of model performance, and the inner loop is used for hyperparameter selection, in order to avoid information leakage during the hyperparameter search process. Specifically, we predefined basic hyperparameter search ranges and employed the Optuna optimization framework to perform intelligent hyperparameter search within these ranges [1]. For each method, 20 hyperparameter optimization trials were conducted, and each trial was evaluated using 10-fold cross-validation, consistent with the training procedure. Detailed hyperparameter configurations are provided in Table S1, Table S2, Table S3.

**Table S1.** Hyperparameter settings and search ranges for feature engineering–based machine learning methods in GPBench.

| Method | Hyperparameters | Ranges |
|---|---|---|
| ElasticNet | *alpha* | [10<sup>−4</sup>, 10<sup>1</sup>] |
|  | *ℓ<sub>1</sub>_ratio* | [0, 1] |
| SVR | *C* | [10<sup>−3</sup>, 1] |
|  | *epsilon* | [0.01, 1] |
|  | *kernel* | {*RBF*, *Polynomial*} |
|  | *gamma* | {*scale*, *auto*} |
|  | *degree* | [1, 5] |
| RF | *n_estimators* | [100, 1000] |
|  | *max_depth* | [3, 10] |
| XGBoost | *learning_rate* | [0.01, 0.2] |
|  | *n_estimators* | [50, 1000] |
|  | *max_depth* | [3, 10] |
|  | *min_child_weight* | [1, 10] |
|  | *subsample* | [0.05, 1.0] |
|  | *colsample_bytree* | [0.5, 1.0] |
|  | *gamma* | [0, 10] |
|  | *reg_alpha* | [10<sup>−3</sup>, 10] |
|  | *reg_lambda* | [10<sup>−3</sup>, 10] |
| LightGBM | *learning_rate* | [10<sup>−3</sup>, 0.2] |
|  | *num_leaves* | [15, 255] |
|  | *min_data_in_leaf* | [10, 100] |
|  | *max_depth* | [3, 10] |
|  | *lambda_l*1 | [0.0, 5.0] |
|  | *lambda_l*2 | [0.0, 5.0] |
|  | *min_gain_to_split* | [0.0, 5.0] |
|  | *feature_fraction* | [0.6, 1.0] |
|  | *bagging_fraction* | [0.6, 1.0] |
|  | *bagging_freq* | [0, 10] |
|  | *num_boost_round* | [100, 1000] |

**Table S2.** Hyperparameter settings and search ranges for CNN-based methods in GPBench.

| Method | Hyperparameters | Ranges |
|---|---|---|
| DeepGS | *learning_rate* | [10<sup>−4</sup>, 0.1] |
|  | *momentum* | [0.1, 0.9] |
|  | *batch_size* | {32, 64, 128} |
|  | *weight_decay* | {10<sup>−4</sup>, 10<sup>−3</sup>, 10<sup>−2</sup>, 10<sup>−1</sup>} |
|  | *patience* | [10, 100] |
| DLGWAS | *learning_rate* | [10<sup>−4</sup>, 0.1] |
|  | *batch_size* | {32, 64, 128} |
|  | *patience* | [5, 10] |
| G2PDeep | *learning_rate* | [10<sup>−4</sup>, 0.1] |
|  | *batch_size* | {32, 64, 128} |
|  | *patience* | [1, 10] |
| MVP | *learning_rate* | [10<sup>−4</sup>, 0.1] |
|  | *batch_size* | {16, 32, 64} |
|  | *patience* | [1, 10] |
|  | *nb_filters* | {8, 16, 32} |
| DNNGP | *learning_rate* | [10<sup>−4</sup>, 0.1] |
|  | *patience* | [10, 100] |
|  | *batch_size* | {32, 64, 128} |
|  | *dropout*1 | [0.0, 0.9] |
|  | *dropout*2 | [0.0, 0.9] |
|  | *weight_decay* | {10<sup>−5</sup>, 10<sup>−4</sup>, 10<sup>−3</sup>} |

**Table S3.** Hyperparameter settings and search ranges for Hybrid-based methods in GPBench.

| Method | Hyperparameters | Ranges |
|---|---|---|
| EIR | *learning_rate* | [10<sup>−4</sup>, 0.1] |
|  | *batch_size* | {32, 64, 128} |
|  | *patience* | [10, 100] |
| SoyDNGP | *learning_rate* | [10<sup>−4</sup>, 0.1] |
|  | *batch_size* | {32, 64, 128} |
|  | *patience* | [1, 10] |
| DeepCCR | *learning_rate* | [10<sup>−4</sup>, 0.1] |
|  | *batch_size* | {32, 64, 128} |
|  | *patience* | [1, 10] |
| Cropformer | *learning_rate* | [10<sup>−4</sup>, 10<sup>−1</sup>] |
|  | *heads* | [1, 8] |
|  | *dropout* | [0.1, 0.9] |
|  | *batch_size* | {32, 64, 128} |
| GEFormer | *learning_rate* | [10<sup>−4</sup>, 0.1] |
|  | *batch_size* | {32, 64, 128} |
|  | *patience* | [1, 10] |
| CropARNet | *learning_rate* | [10<sup>−4</sup>, 0.1] |
|  | *batch_size* | {32, 64, 128} |
|  | *weight_decay* | {10<sup>−4</sup>, 10<sup>−3</sup>, 10<sup>−2</sup>, 10<sup>−1</sup>} |
|  | *patience* | [1, 10] |

## B Additional Experimental Results

### B.1 Supplementary Effectiveness Analysis

#### B.1.1 Multi-species regression tasks

In this section, we further report additional evaluation metrics for regression tasks across multiple species, including MAE (Table S1), RMSE (Table S2), and *R*^2^ (Table S3). Based on the MAE results, substantial performance differences among models were observed across different species. From the perspective of model categories, genetic-statistics-based models, including rrBLUP, GBLUP, and BayesA/B/C, showed relatively small MAE variation across most species, with low standard deviations (predominantly in the green or yellow ranges), demonstrating strong stability and robustness. Overall, these methods rarely suffered from extreme performance degradation, making them reliable and strong baseline models. Feature-engineeringbased machine learning models also achieved low MAE on most species and exhibited balanced performance, with overall accuracy comparable to or even exceeding that of genetic-statistics-based models.

Among convolution-based deep learning models, G2PDeep and DLGWAS showed similar performance patterns, likely due to their shared dual-channel architectures; however, their overall performance did not significantly outperform genetic-statistics-based approaches. Other convolutional neural networks exhibited pronounced instability and species dependency. For example, DNNGP produced substantially higher MAE on multiple species, indicating high sensitivity to data scale and noise. Hybrid architecture-based models demonstrated clearly divergent performance characteristics. EIR achieved MAE values comparable to those of traditional genetic-statistics-based models across most species, whereas SoyDNGP and Cropformer exhibited extremely large errors on multiple datasets, reflecting poor generalization ability. Consistent patterns were also observed for RMSE and *R*^2^, supporting the conclusions drawn from the MAE analysis.

#### B.1.2 Multi-species classification tasks

In addition to conventional regression tasks, we systematically evaluated model performance on classification tasks. Specifically, experiments were conducted on a horse coat color trait dataset and a human age-related macular degeneration (AMD) dataset. The coat color prediction task is a multi-class classification problem, whereas the AMD prediction task is a binary classification problem. The experimental results are shown in Figure S2.

For the multi-class classification task, clear performance differences were observed among different models. Overall, consistent with their performance in regression tasks, ensemble learning and tree-based models (XGBoost and LightGBM) achieved the best performance across all four evaluation metrics. Genetic statistics-based models (BayesA/B and rrBLUP) remained relatively stable. In contrast, CNN-based models exhibited smaller intra-model variability, whereas hybrid-architecture deep learning models showed substantial performance variation, with EIR approaching or even surpassing the genetic-statistics-based methods.

For the binary classification task, although the models exhibited similar performance in terms of accuracy and recall, significant differences were observed in precision and F1-score, indicating a severe class imbalance in the dataset. Consequently, precision and F1-score provide a more reliable evaluation of model performance. Notably, model performance under these two metrics follows trends consistent with those observed in the multi-class classification task.

#### B.1.3 Genetic effect analysis

In Section 4.1.2, we analyzed three traits in the Cattle dataset and found that they exhibit distinct genetic architectures, as shown in Fig. S3. By calculating the P-values between genotypes and phenotypes, clear signal peaks can be observed in the original features of the mkg and fpro traits, with the fpro trait showing more pronounced peaks. In contrast, no evident peak is observed for the scs trait.

Figure S4 illustrates the performance of 21 GP models. The experimental results indicate that the three traits—fpro, mkg, and scs—exhibit a clear high–medium–low trend in terms of genetic architecture complexity.

For the fpro trait, removing major-effect loci leads to a significant decline in predictive performance for most models. When predictions are conducted using only major-effect loci, some models fail to recover their original performance. In contrast, models such as RF, XGBoost, and G2PDeep are able to recover, and in some cases even surpass, the original predictive performance. This suggests that for traits jointly influenced by major-effect loci and a large number of minor-effect loci, existing models tend to focus on exploiting SNPs with pronounced effects, while their ability to capture minor-effect loci is highly imbalanced due to differences in model architectures and learning mechanisms.

For the mkg trait, the loci removed correspond to moderate-effect loci, and their removal similarly results in a significant decrease in predictive performance. When predictions are performed using only the removed moderate-effect loci, most models struggle to recover their original performance, and their predictive accuracy generally does not exceed that achieved by the collective contribution of minor-effect loci. This indicates that for traits primarily governed by moderate-effect loci together with numerous minor-effect loci, model performance depends not only on the strength of individual loci but also on the ability to jointly model weak signals across multiple loci, an aspect in which current models remain limited.

For the scs trait, the loci removed correspond to minor-effect loci, and the predictive performance of most models does not show a significant decline after their removal. However, when predictions rely solely on the removed minor-effect loci, model performance decreases markedly. This implies that for traits mainly controlled by a large number of minor-effect loci, localized weak-effect signals are insufficient to support effective prediction, and the genetic information is instead embedded in the cumulative effects of many weakly contributing loci.

### B.2 Supplementary Sensitivity Analysis

We systematically evaluated the sensitivity of different models to variations in sample size, feature dimensionality, and random seed initialization. Figure S5 illustrates the responses of 21 models under different sample sizes and random initialization settings; a detailed analysis of these experimental results is provided in Section 4.2 of the main text. Figure S6 also presents the responses of all models under varying feature dimensionalities and random initialization conditions.

The experimental results show that genetic statistical methods (e.g., rrBLUP, GBLUP, and BayesA/B/C) as well as feature-engineered classical machine learning methods (e.g., ElasticNet and SVR) exhibit good stability across different feature dimensions and random initializations, with relatively small performance fluctuations. Tree-based ensemble methods (RF, XGBoost, and LightGBM) not only demonstrate remarkable stability but also consistently achieve high predictive performance. In contrast, CNN-based deep learning methods display pronounced intra-model variability under different initialization settings. For example, DeepGS, G2PDeep, and MVP show relatively stable behavior, whereas DLGWAS exhibits large performance variance, and DNNGP even fails to converge in some cases. Similarly, hybrid architectures-based methods (e.g., SoyDNGP and Cropformer) present more substantial performance fluctuations. Overall, these results indicate that deep learning methods are generally more sensitive to input feature dimensionality and random initialization than genetic statistical and traditional machine learning methods.

### B.3 Supplementary Robustness Analysis

In Section 4.3, we analyze the impact of random noise on the PCC performance of different models for the Yeast trait (*P*/*N* = 11.53, normal distributions) and the Rice grain weight trait (*P*/*N* = 2.53, non-normal distributions). Figure S7 illustrates the effects of random noise on the *R*^2^ of model performance, and the results further support the conclusions presented in the main text.

**Table S4.** Mean Absolute Error of Benchmark Methods on Various Species.

| Methods | Cattle | Chicken | Chickpea | Cotton | Loblolly pine | Maize | Millet | Mouse | Pig | Rapeseed | Rice | Soybean | Wheat | Yeast |
|---|---|---|---|---|---|---|---|---|---|---|---|---|---|---|
| rrBLUP | 0.496 | 2.500 | 5.531 | 1.121 | 0.882 | 1.707 | 6.020 | 0.363 | 8.774 | 4.106 | 0.153 | 5.151 | 361.6 | 1.636 |
| GBLUP | 0.512 | 2.529 | 5.544 | 1.127 | 0.892 | 1.709 | 6.063 | 0.365 | 8.783 | 4.097 | 0.154 | 5.161 | 365.0 | 1.637 |
| BayesA | 0.484 | 2.494 | 5.562 | 1.117 | 0.886 | 1.711 | 6.000 | 0.364 | 8.812 | 4.005 | 0.154 | 5.148 | 359.5 | 1.609 |
| BayesB | 0.504 | 2.559 | 5.878 | 1.132 | 0.890 | 1.762 | 6.188 | 0.400 | 9.272 | 5.209 | 0.172 | 5.315 | 366.1 | 1.608 |
| BayesC | 0.550 | 2.544 | 5.889 | 1.121 | 0.887 | 1.796 | 6.211 | 0.376 | 8.993 | 6.115 | 0.159 | 5.281 | 365.3 | 1.649 |
| ElasticNet | 0.497 | 2.497 | 5.508 | 1.118 | 0.897 | 1.739 | 5.950 | 0.366 | 8.735 | 4.041 | 0.156 | 5.228 | 361.9 | 1.598 |
| SVR | 0.500 | 2.530 | 5.508 | 1.039 | 0.877 | 1.730 | 5.894 | 0.362 | 8.705 | 3.861 | 0.153 | 4.927 | 398.2 | 1.668 |
| RF | 0.707 | 2.649 | 5.572 | 1.051 | 0.855 | 1.863 | 6.286 | 0.369 | 9.717 | 4.350 | 0.149 | 4.972 | 368.8 | 1.773 |
| XGBoost | 0.574 | 2.563 | 5.542 | 1.006 | 0.866 | 1.801 | 5.632 | 0.367 | 8.647 | 3.885 | 0.156 | 4.564 | 356.1 | 1.634 |
| LightGBM | 0.635 | 2.561 | 6.833 | 1.033 | 0.850 | 1.786 | 5.937 | 0.367 | 9.732 | 3.962 | 0.164 | 4.634 | 373.4 | 1.749 |
| DeepGS | 0.548 | 5.528 | 13.02 | 2.472 | 0.888 | 6.699 | 6.092 | 0.374 | 16.68 | 16.51 | 0.206 | 5.843 | 645.7 | 1.644 |
| DLGWAS | 0.567 | 2.858 | 5.615 | 1.561 | 0.846 | 2.095 | 6.267 | 0.366 | 9.217 | 5.033 | 0.299 | 5.296 | 377.3 | 31.53 |
| G2PDeep | 0.574 | 2.657 | 5.614 | 1.270 | 0.853 | 1.801 | 6.156 | 0.368 | 8.880 | 4.133 | 0.158 | 5.263 | 364.7 | 1.687 |
| MVP | 0.813 | 2.556 | 6.192 | 1.198 | 0.989 | 1.822 | 6.704 | 0.500 | 17.78 | 4.165 | 0.185 | 5.722 | 371.4 | 1.740 |
| DNNGP | 0.582 | 11.98 | 7.559 | 9.351 | 0.933 | 35.87 | 8.015 | 0.398 | 10.75 | 15.72 | 0.322 | 7.647 | 780.7 | 1.963 |
| EIR | 0.529 | 3.007 | 5.842 | 1.126 | 0.988 | 1.958 | 6.212 | 0.397 | 10.43 | 4.239 | 0.157 | 5.117 | 370.2 | 1.641 |
| SoyDNGP | 1.484 | 3.417 | 6.571 | 1.178 | 0.973 | 6.091 | 8.689 | 0.423 | 12.06 | 37.09 | 0.352 | 7.030 | 414.9 | 2.007 |
| DeepCCR | 0.829 | 2.704 | 5.670 | 1.269 | 0.928 | 2.247 | 7.965 | 0.394 | 10.51 | 5.060 | 0.155 | 5.617 | 371.5 | 2.017 |
| Cropformer | 0.743 | 11.47 | 9.503 | 1.551 | 1.032 | 22.80 | 8.654 | 0.505 | 52.83 | 35.98 | 0.871 | 22.72 | 417.4 | 1.802 |
| GEFormer | 0.572 | 3.277 | 5.674 | 1.540 | 0.929 | 3.283 | 6.566 | 0.389 | 9.538 | 8.377 | 0.203 | 5.578 | 420.7 | 1.750 |
| CropARNet | 0.549 | 2.785 | 5.563 | 1.162 | 0.971 | 2.210 | 6.412 | 0.379 | 9.857 | 5.689 | 0.161 | 5.235 | 374.2 | 1.732 |

Color legend: (*std* ≤ 0.05) (0.05 < *std* ≤ 0.1) (0.1 < *std* ≤ 0.5) (0.5 < *std* ≤ 1) (*std* > 1)

**Table S5.**
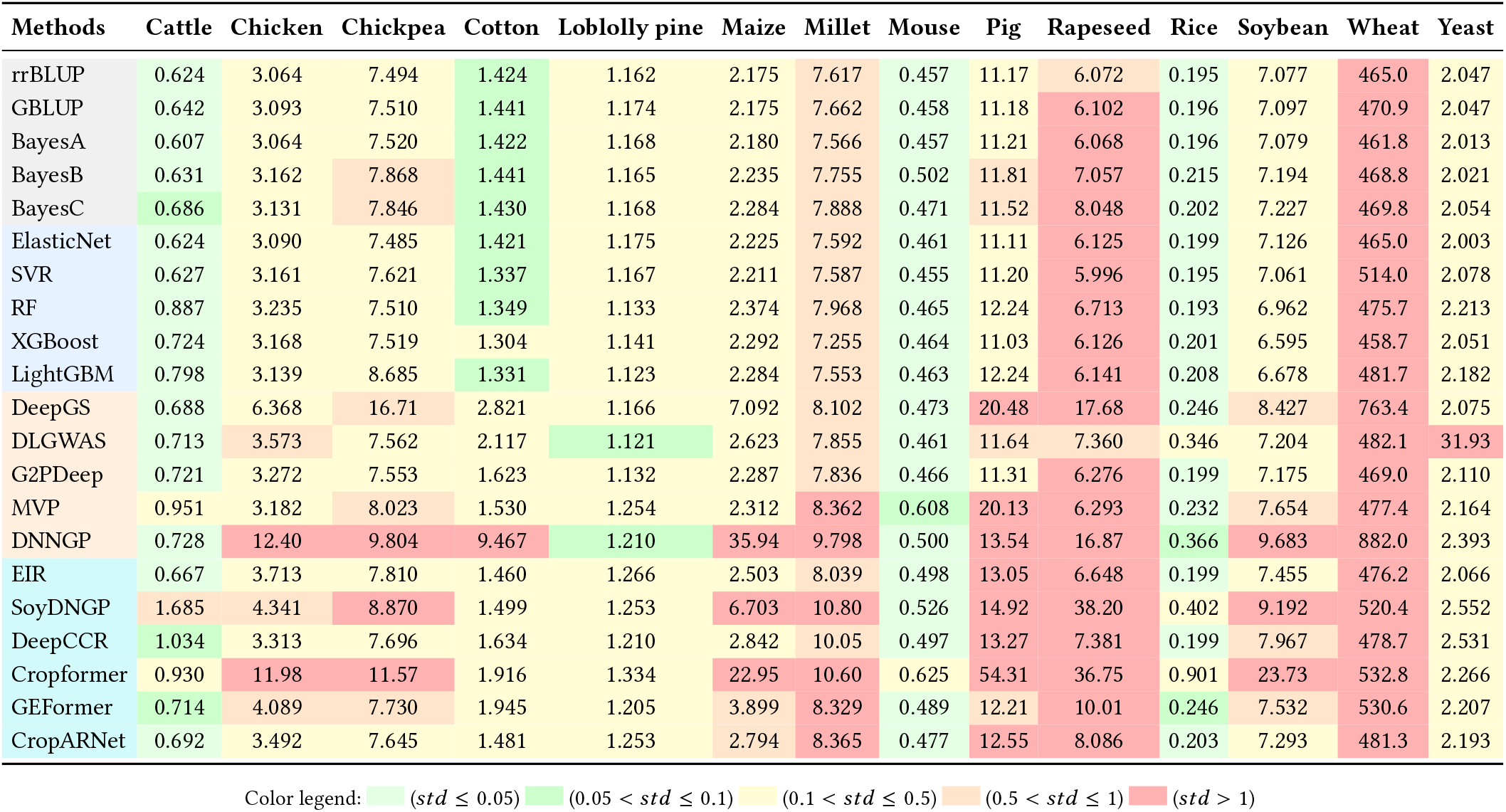
Root Mean Squared Error of Benchmark Methods on Various Species.

| Methods | Cattle | Chicken | Chickpea | Cotton | Loblolly pine | Maize | Millet | Mouse | Pig | Rapeseed | Rice | Soybean | Wheat | Yeast |
|---|---|---|---|---|---|---|---|---|---|---|---|---|---|---|
| rrBLUP | 0.624 | 3.064 | 7.494 | 1.424 | 1.162 | 2.175 | 7.617 | 0.457 | 11.17 | 6.072 | 0.195 | 7.077 | 465.0 | 2.047 |
| GBLUP | 0.642 | 3.093 | 7.510 | 1.441 | 1.174 | 2.175 | 7.662 | 0.458 | 11.18 | 6.102 | 0.196 | 7.097 | 470.9 | 2.047 |
| BayesA | 0.607 | 3.064 | 7.520 | 1.422 | 1.168 | 2.180 | 7.566 | 0.457 | 11.21 | 6.068 | 0.196 | 7.079 | 461.8 | 2.013 |
| BayesB | 0.631 | 3.162 | 7.868 | 1.441 | 1.165 | 2.235 | 7.755 | 0.502 | 11.81 | 7.057 | 0.215 | 7.194 | 468.8 | 2.021 |
| BayesC | 0.686 | 3.131 | 7.846 | 1.430 | 1.168 | 2.284 | 7.888 | 0.471 | 11.52 | 8.048 | 0.202 | 7.227 | 469.8 | 2.054 |
| ElasticNet | 0.624 | 3.090 | 7.485 | 1.421 | 1.175 | 2.225 | 7.592 | 0.461 | 11.11 | 6.125 | 0.199 | 7.126 | 465.0 | 2.003 |
| SVR | 0.627 | 3.161 | 7.621 | 1.337 | 1.167 | 2.211 | 7.587 | 0.455 | 11.20 | 5.996 | 0.195 | 7.061 | 514.0 | 2.078 |
| RF | 0.887 | 3.235 | 7.510 | 1.349 | 1.133 | 2.374 | 7.968 | 0.465 | 12.24 | 6.713 | 0.193 | 6.962 | 475.7 | 2.213 |
| XGBoost | 0.724 | 3.168 | 7.519 | 1.304 | 1.141 | 2.292 | 7.255 | 0.464 | 11.03 | 6.126 | 0.201 | 6.595 | 458.7 | 2.051 |
| LightGBM | 0.798 | 3.139 | 8.685 | 1.331 | 1.123 | 2.284 | 7.553 | 0.463 | 12.24 | 6.141 | 0.208 | 6.678 | 481.7 | 2.182 |
| DeepGS | 0.688 | 6.368 | 16.71 | 2.821 | 1.166 | 7.092 | 8.102 | 0.473 | 20.48 | 17.68 | 0.246 | 8.427 | 763.4 | 2.075 |
| DLGWAS | 0.713 | 3.573 | 7.562 | 2.117 | 1.121 | 2.623 | 7.855 | 0.461 | 11.64 | 7.360 | 0.346 | 7.204 | 482.1 | 31.93 |
| G2PDeep | 0.721 | 3.272 | 7.553 | 1.623 | 1.132 | 2.287 | 7.836 | 0.466 | 11.31 | 6.276 | 0.199 | 7.175 | 469.0 | 2.110 |
| MVP | 0.951 | 3.182 | 8.023 | 1.530 | 1.254 | 2.312 | 8.362 | 0.608 | 20.13 | 6.293 | 0.232 | 7.654 | 477.4 | 2.164 |
| DNNGP | 0.728 | 12.40 | 9.804 | 9.467 | 1.210 | 35.94 | 9.798 | 0.500 | 13.54 | 16.87 | 0.366 | 9.683 | 882.0 | 2.393 |
| EIR | 0.667 | 3.713 | 7.810 | 1.460 | 1.266 | 2.503 | 8.039 | 0.498 | 13.05 | 6.648 | 0.199 | 7.455 | 476.2 | 2.066 |
| SoyDNGP | 1.685 | 4.341 | 8.870 | 1.499 | 1.253 | 6.703 | 10.80 | 0.526 | 14.92 | 38.20 | 0.402 | 9.192 | 520.4 | 2.552 |
| DeepCCR | 1.034 | 3.313 | 7.696 | 1.634 | 1.210 | 2.842 | 10.05 | 0.497 | 13.27 | 7.381 | 0.199 | 7.967 | 478.7 | 2.531 |
| Cropformer | 0.930 | 11.98 | 11.57 | 1.916 | 1.334 | 22.95 | 10.60 | 0.625 | 54.31 | 36.75 | 0.901 | 23.73 | 532.8 | 2.266 |
| GEFormer | 0.714 | 4.089 | 7.730 | 1.945 | 1.205 | 3.899 | 8.329 | 0.489 | 12.21 | 10.01 | 0.246 | 7.532 | 530.6 | 2.207 |
| CropARNet | 0.692 | 3.492 | 7.645 | 1.481 | 1.253 | 2.794 | 8.365 | 0.477 | 12.55 | 8.086 | 0.203 | 7.293 | 481.3 | 2.193 |

Color legend: (*std* ≤ 0.05) (0.05 < *std* ≤ 0.1) (0.1 < *std* ≤ 0.5) (0.5 < *std* ≤ 1) (*std* > 1)

**Table S6.** Coefficient of Determination of Benchmark Methods on Various Species.

| **Methods** | **Cattle** | **Chicken** | **Chickpea** | **Cotton** | **Loblolly pine** | **Maize** | **Millet** | **Mouse** | **Pig** | **Rapeseed** | **Rice** | **Soybean** | **Wheat** | **Yeast** |
|---|---|---|---|---|---|---|---|---|---|---|---|---|---|---|
| rrBLUP | 0.608 | 0.113 | 0.393 | 0.359 | 0.066 | 0.413 | 0.411 | 0.155 | 0.473 | 0.655 | 0.653 | 0.362 | 0.238 | 0.455 |
| GBLUP | 0.584 | 0.097 | 0.390 | 0.342 | 0.048 | 0.414 | 0.404 | 0.152 | 0.473 | 0.652 | 0.650 | 0.359 | 0.218 | 0.455 |
| BayesA | 0.629 | 0.113 | 0.389 | 0.361 | 0.055 | 0.412 | 0.418 | 0.154 | 0.470 | 0.656 | 0.649 | 0.362 | 0.248 | 0.473 |
| BayesB | 0.598 | 0.055 | 0.328 | 0.344 | 0.060 | 0.380 | 0.386 | -0.032 | 0.407 | 0.522 | 0.581 | 0.341 | 0.225 | 0.470 |
| BayesC | 0.516 | 0.070 | 0.335 | 0.354 | 0.055 | 0.353 | 0.362 | 0.100 | 0.441 | 0.376 | 0.630 | 0.334 | 0.222 | 0.453 |
| ElasticNet | 0.607 | 0.096 | 0.394 | 0.361 | 0.048 | 0.385 | 0.414 | 0.139 | 0.480 | 0.651 | 0.641 | 0.353 | 0.238 | 0.480 |
| SVR | 0.603 | 0.053 | 0.373 | 0.431 | 0.061 | 0.395 | 0.414 | 0.163 | 0.471 | 0.665 | 0.655 | 0.365 | 0.069 | 0.440 |
| RF | 0.208 | 0.014 | 0.391 | 0.424 | 0.113 | 0.304 | 0.356 | 0.125 | 0.369 | 0.579 | 0.660 | 0.383 | 0.202 | 0.369 |
| XGBoost | 0.473 | 0.051 | 0.389 | 0.455 | 0.100 | 0.350 | 0.465 | 0.129 | 0.487 | 0.649 | 0.632 | 0.445 | 0.258 | 0.455 |
| LightGBM | 0.358 | 0.067 | 0.186 | 0.438 | 0.129 | 0.354 | 0.420 | 0.128 | 0.368 | 0.646 | 0.610 | 0.433 | 0.182 | 0.387 |
| DeepGS | 0.523 | -2.843 | -0.071 | -1.547 | 0.061 | -5.309 | 0.332 | 0.093 | -0.792 | -2.089 | 0.437 | 0.086 | -1.059 | 0.442 |
| DLGWAS | 0.488 | -0.208 | 0.382 | -0.403 | 0.130 | 0.151 | 0.373 | 0.141 | 0.426 | 0.492 | -0.138 | 0.338 | 0.179 | -737.379 |
| G2PDeep | 0.474 | -0.013 | 0.384 | 0.183 | 0.114 | 0.354 | 0.377 | 0.119 | 0.461 | 0.632 | 0.638 | 0.345 | 0.224 | 0.419 |
| MVP | -0.190 | 0.040 | 0.301 | 0.239 | -0.132 | 0.334 | 0.283 | -0.559 | -3.029 | 0.630 | 0.510 | 0.248 | 0.196 | 0.388 |
| DNNGP | 0.466 | -13.795 | -0.056 | -28.177 | -0.018 | -160.953 | -0.011 | -0.012 | 0.215 | -2.599 | -0.267 | -0.238 | -1.992 | 0.262 |
| EIR | 0.552 | -0.310 | 0.339 | 0.324 | -0.109 | 0.225 | 0.341 | -0.013 | 0.277 | 0.581 | 0.642 | 0.294 | 0.200 | 0.444 |
| SoyDNGP | -2.482 | -0.793 | 0.141 | 0.267 | -0.091 | -5.844 | -0.199 | -0.121 | 0.037 | -18.628 | -0.958 | -0.088 | 0.041 | 0.157 |
| DeepCCR | -0.087 | -0.033 | 0.361 | 0.163 | -0.010 | 0.004 | -0.025 | 0.002 | 0.246 | 0.470 | 0.639 | 0.192 | 0.192 | 0.175 |
| Cropformer | 0.111 | -13.474 | -0.481 | -0.218 | -0.266 | -65.078 | -0.152 | -0.676 | -11.479 | -12.278 | -7.015 | -6.410 | -0.049 | 0.321 |
| GEFormer | 0.482 | -0.685 | 0.354 | -0.225 | -0.007 | -0.964 | 0.291 | 0.024 | 0.369 | 0.001 | 0.431 | 0.279 | 0.000 | 0.359 |
| CropARNet | 0.516 | -0.163 | 0.367 | 0.307 | -0.088 | 0.002 | 0.284 | 0.076 | 0.312 | 0.346 | 0.626 | 0.323 | 0.184 | 0.379 |

Color legend: (*std* ≤ 0.05) (0.05 < *std* ≤ 0.1) (0.1 < *std* ≤ 0.5) (0.5 < *std* ≤ 1) (*std* > 1)

**Figure S2.**
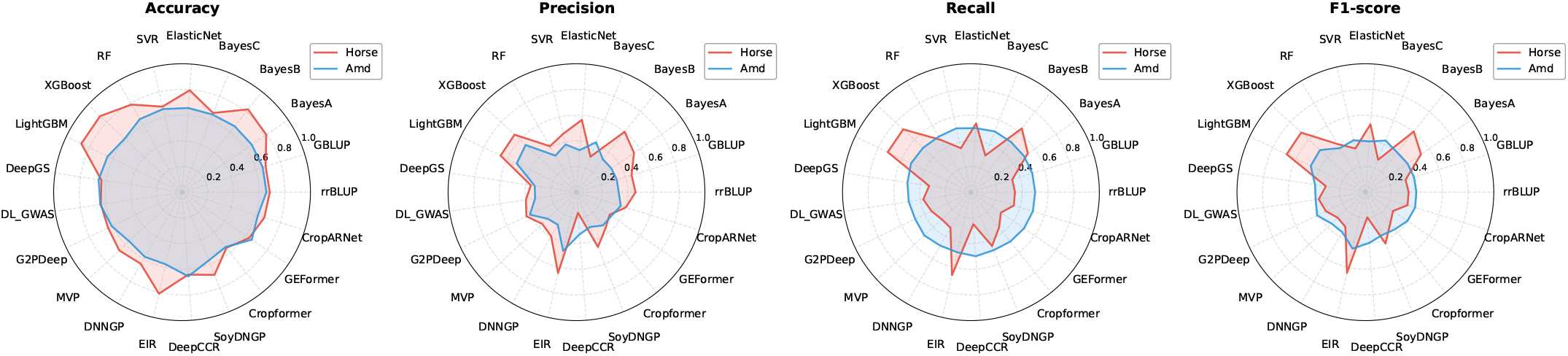
Performance of benchmark methods on classification tasks, showing results on the horse coat color dataset and the human AMD dataset.

To further investigate how models behave under different types of noise, we evaluated the effect of missing on the Pig back fat thickness trait (*P*/*N* = 7.82, normal distributions) and the Rapeseed Flower Time trait (*P*/*N* = 10.03, non-normal distributions), with missing intensities set at 0.1, 0.3, and 0.5. The experimental results are shown in Figure S8. The results indicate that genetic statistical methods (such as rrBLUP and GBLUP) are relatively robust under missing conditions. This is primarily because these models are based on the core assumption of “true signal + normally distributed random noise,” treating noise as an inherent component rather than “useful information” that must be fully fitted. Consequently, they are less prone to overfitting when faced with missing or random noise. Moreover, combining these methods with regularization techniques (such as ridge regression and LASSO) can further mitigate parameter estimation instability caused by noise. In contrast, as missing intensity increases, feature-engineered classical machine learning methods, CNN-based deep learning methods, and hybrid-architecture deep learning models all exhibit pronounced performance fluctuations and a clear downward trend. Furthermore, under higher missing levels, the performance distributions become increasingly dispersed, indicating greater sensitivity to data missingness. This is because missing severely distorts the true data distribution, thereby significantly interfering with the models’ ability to learn underlying genotype–phenotype associations.

**Figure S3.**
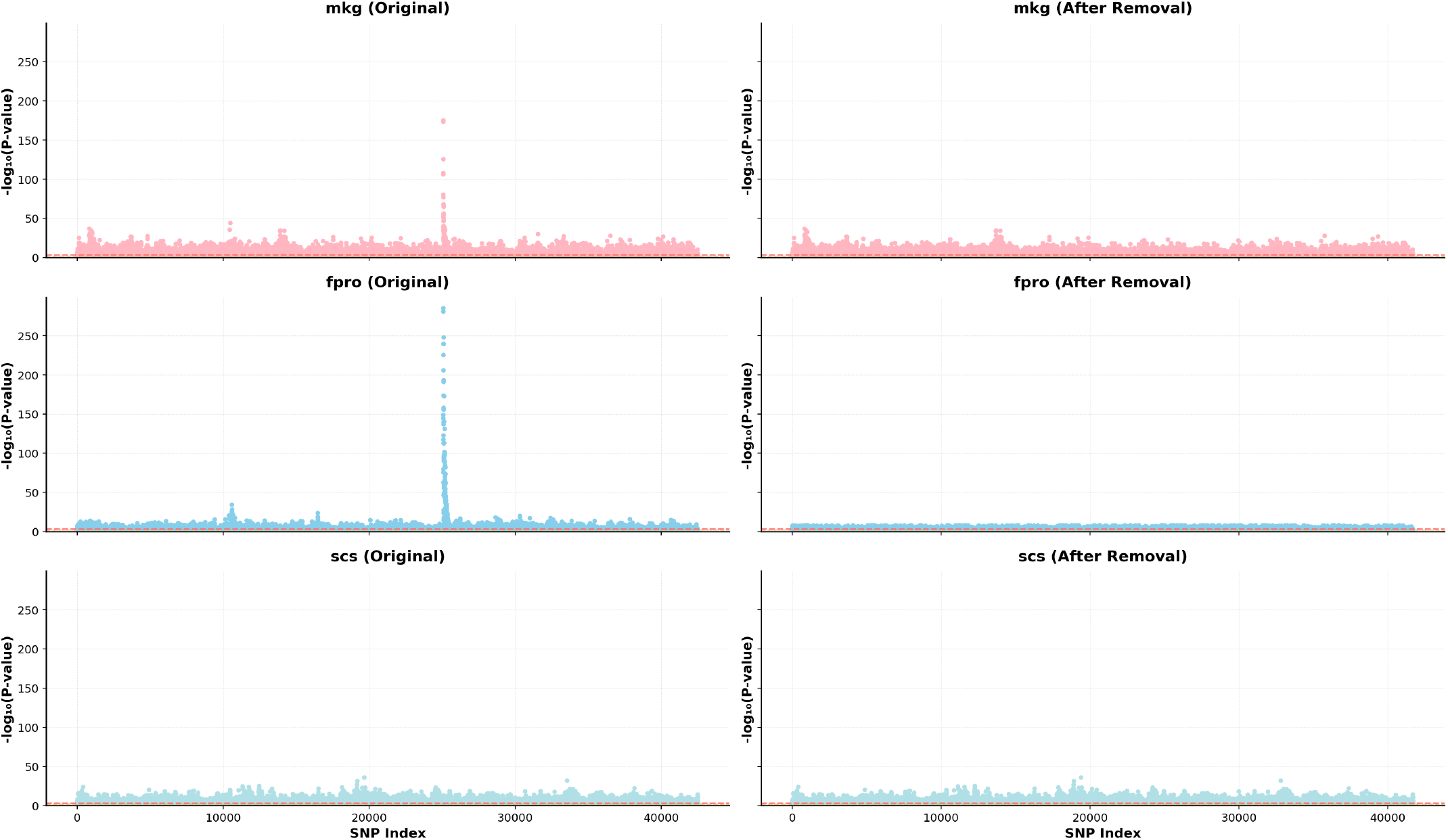
The effect sizes of mkg, fpro, and scs traits in the cattle dataset, which were obtained by calculating the P-values between genotypes and phenotypes.

**Figure S4.**
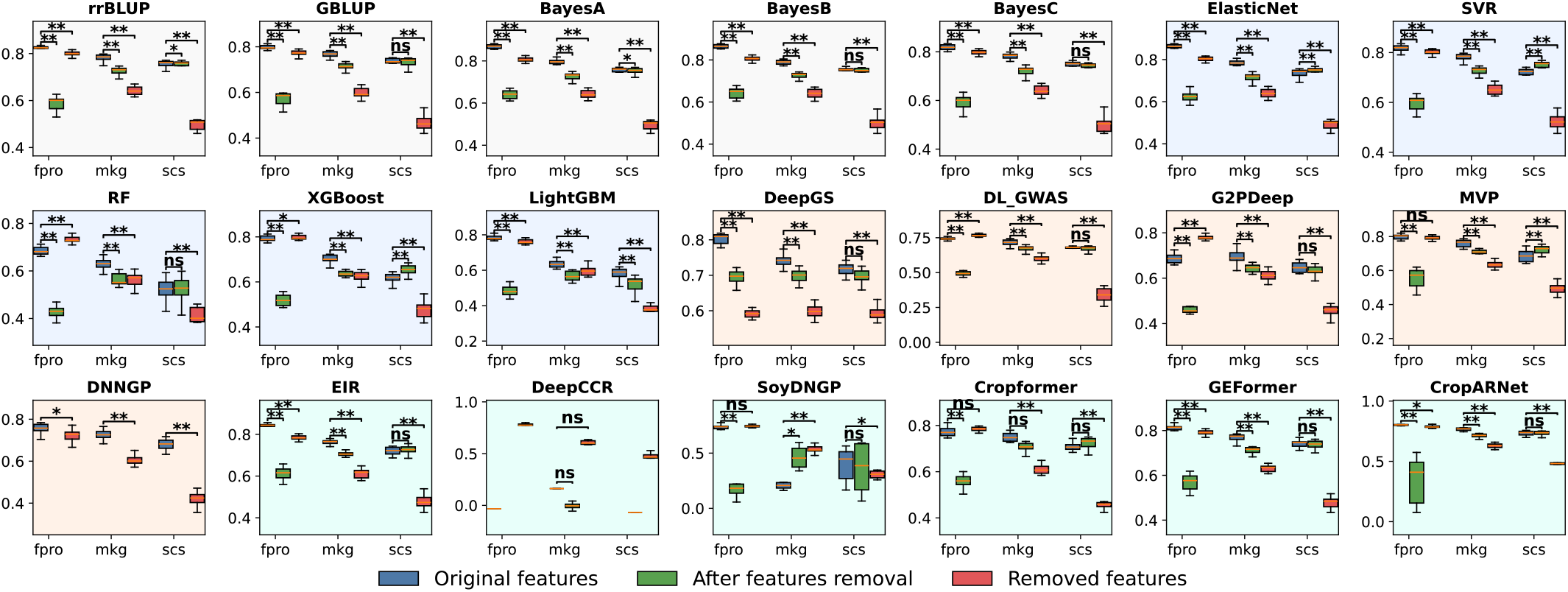
shows the performance differences of 21 GP methods across traits with different genetic architectures. ** indicates *p <* 0.01, * indicates *p <* 0.05, and *ns* indicates not significant.

### B.4 Online Application Platform

To facilitate the use and analysis of existing GP methods by the research community, we developed the GPBench application platform, which comprises multiple modules, including tools, analysis, agent, and task. Detailed information is available in the help section of https://www.sdu-idea.cn/GPBench/. In the following, we focus on the Agent module.

**Figure S5.**
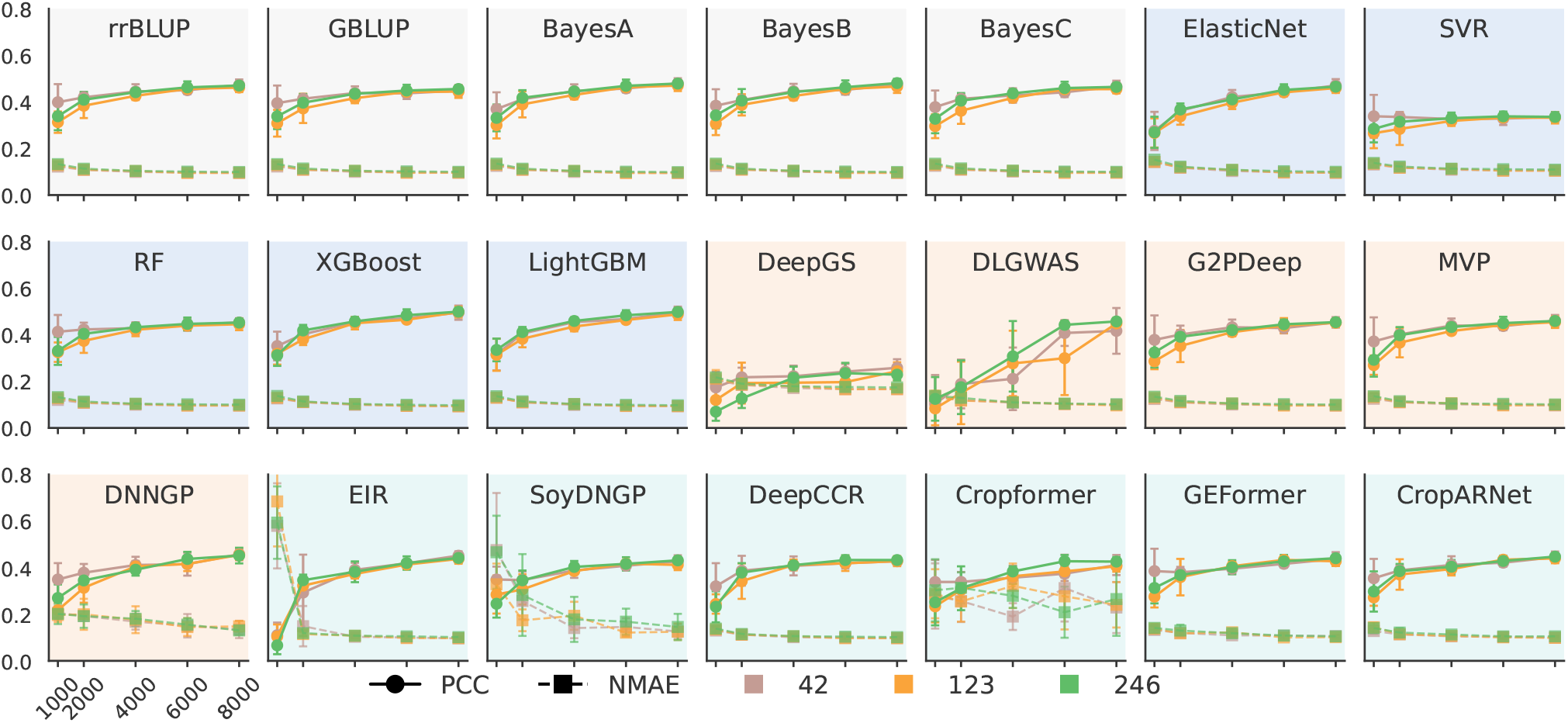
shows the performance of 21 GP methods under different sample sizes and random seed splits, where circles represent PCC, squares represent NMAE, and different colors indicate different random seeds.

**Figure S6.**
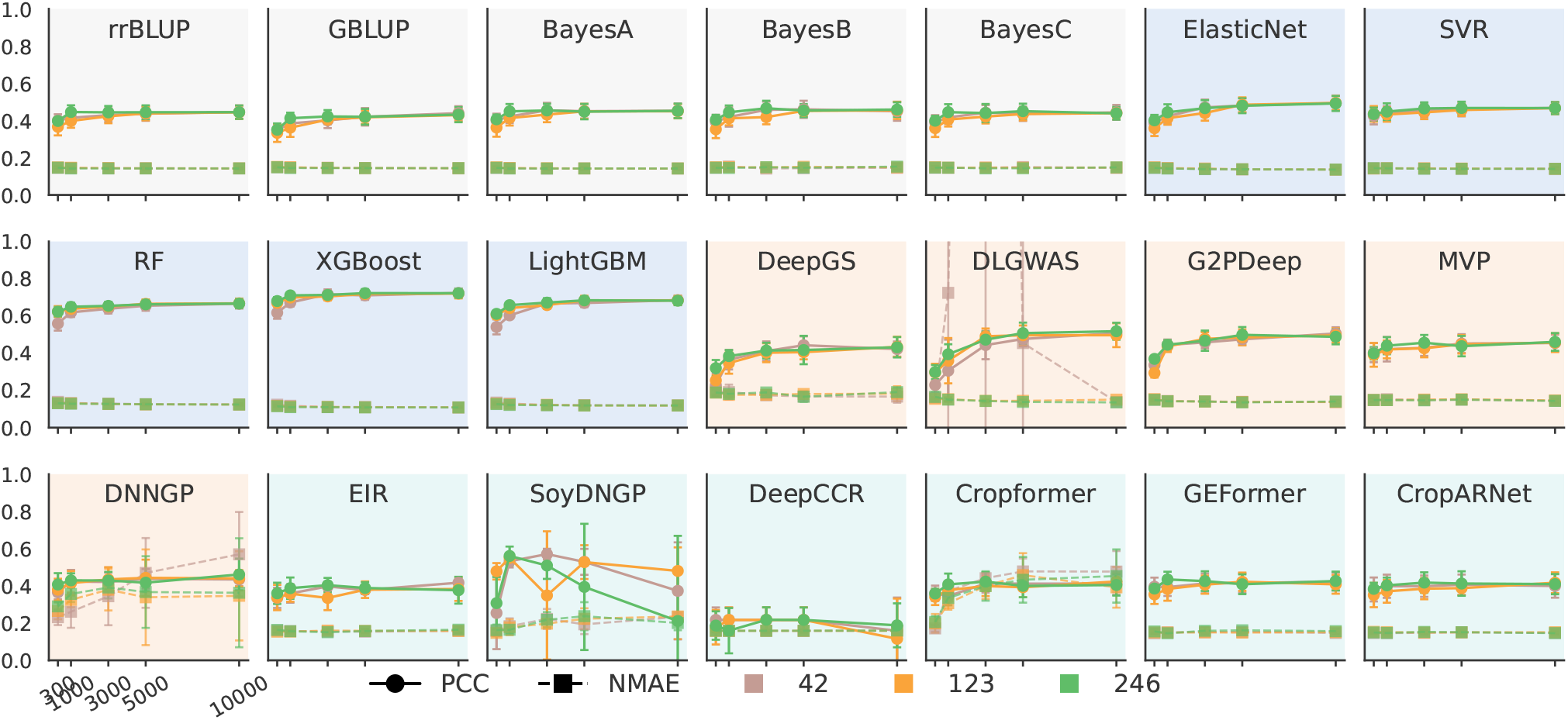
shows the performance of 21 GP methods under different feature dimensions and random seed splits, where circles represent PCC, squares represent NMAE, and different colors indicate different random seeds.

To enable effective interpretation of large-scale benchmark results, we developed an agent-based intelligent data analysis tool, termed the GPBench Agent, supported by the Qwen-Plus large language model.

#### (i) Result analysis task case study

When users choose not to provide a dataset, they can ask questions directly, for example: “What patterns do the models exhibit for the maize DTT trait, and what insights can be derived?” GPBench Agent will then summarize the existing patterns based on the performance of 21 models across 16 species and 57 traits. The agent provides specific answers on how each model performs, the corresponding evaluation metrics, and derives insightful patterns from the aggregated results, see Figure S9.

**Figure S7.**
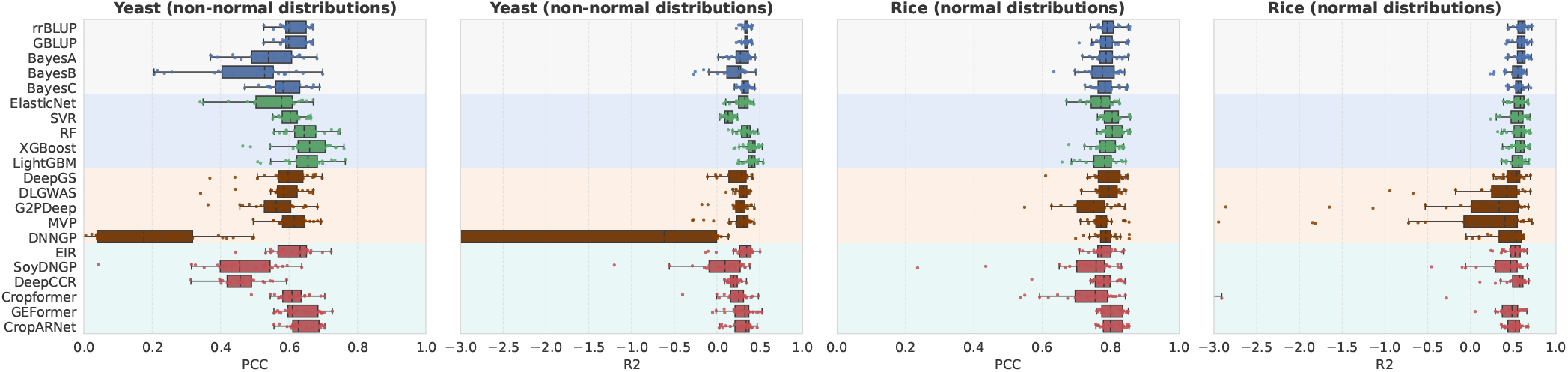
The boxplots illustrate the distributions of PCC and R2 for 21 GP algorithms under different random noise levels. A larger spread of the boxplots indicates greater performance variability under random noise perturbations, reflecting the weaker robustness of the models.

**Figure S8.**
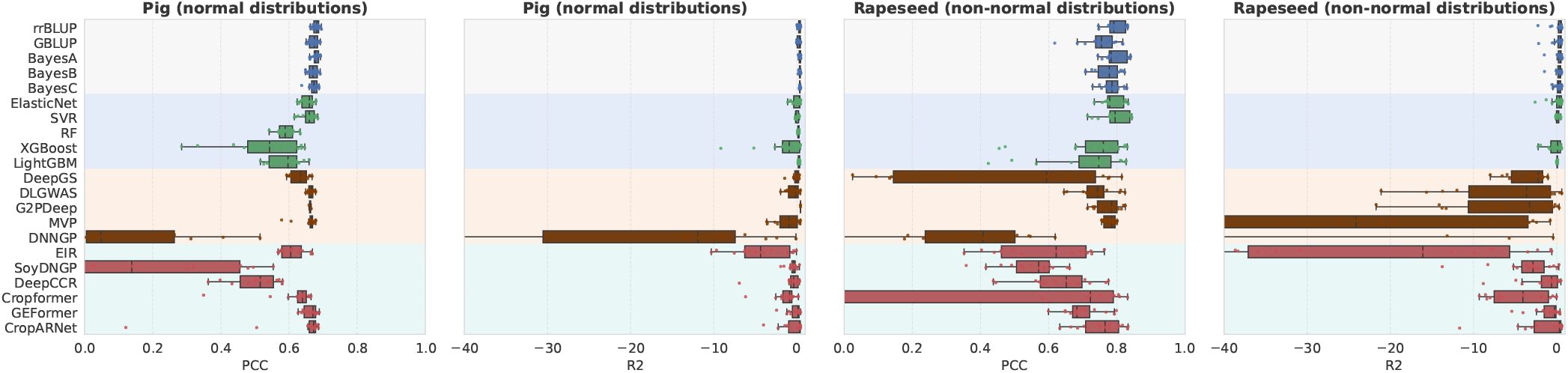
The boxplots illustrate the distributions of PCC and R2 for 21 GP algorithms under different missing levels. A larger spread of the boxplots indicates greater performance variability under missing perturbations, reflecting the weaker robustness of the models.

#### (ii) Algorithm recommendation task case study

When users choose to provide a dataset, they can ask questions directly, for example: “Could you recommend the most suitable algorithm for this dataset?” GPBench Agent first interprets the user’s requirements and analyzes the characteristics of the dataset. The agent then identifies the most similar results from existing historical benchmark data. For instance, it may return: “Found 5 similar datasets: Wheat/Adj. Grain Yield, Pig/Age, Rapeseed/Flowering Time, Millet/MSH, Millet/SGN.” Based on the performance of 21 models on these datasets, the agent provides a recommendation: “XGBoost, Confidence Score: 95%. The upstream answer explicitly recommends XGBoost as the most suitable algorithm for the dataset, based on its superior performance in accuracy, robustness, computational efficiency, memory usage, and generalizability.”

## C GPBench Platform Code and Dataset

This study opens sources and provides the following resources for the community.

**Dataset repository**: https://huggingface.co/datasets/xw97/GPBench.

**GPBench Platform**: https://www.sdu-idea.cn/GPBench/.

**Code repository**: https://github.com/xwzhang2118/GPBench.

**Python library** https://pypi.org/project/gpbench/

Permission to make digital or hard copies of all or part of this work for personal or classroom use is granted without fee provided that copies are not made or distributed for profit or commercial advantage and that copies bear this notice and the full citation on the first page. Copyrights for components of this work owned by others than the author(s) must be honored. Abstracting with credit is permitted. To copy otherwise, or republish, to post on servers or to redistribute to lists, requires prior specific permission and/or a fee. Request permissions from.

**Figure S9.**
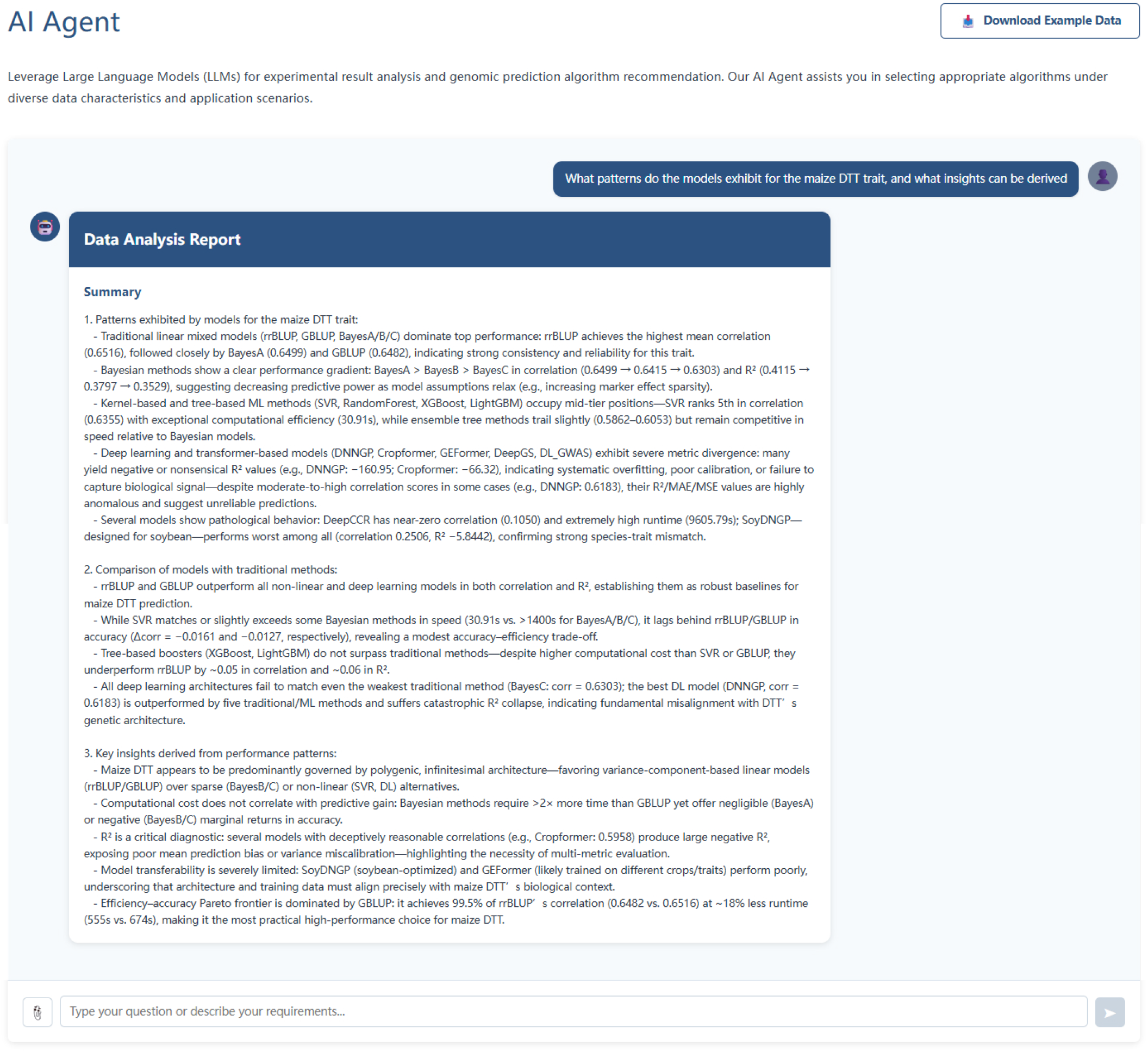
shows the user and agent chat process in data analysis tasks.

**Figure S10.**
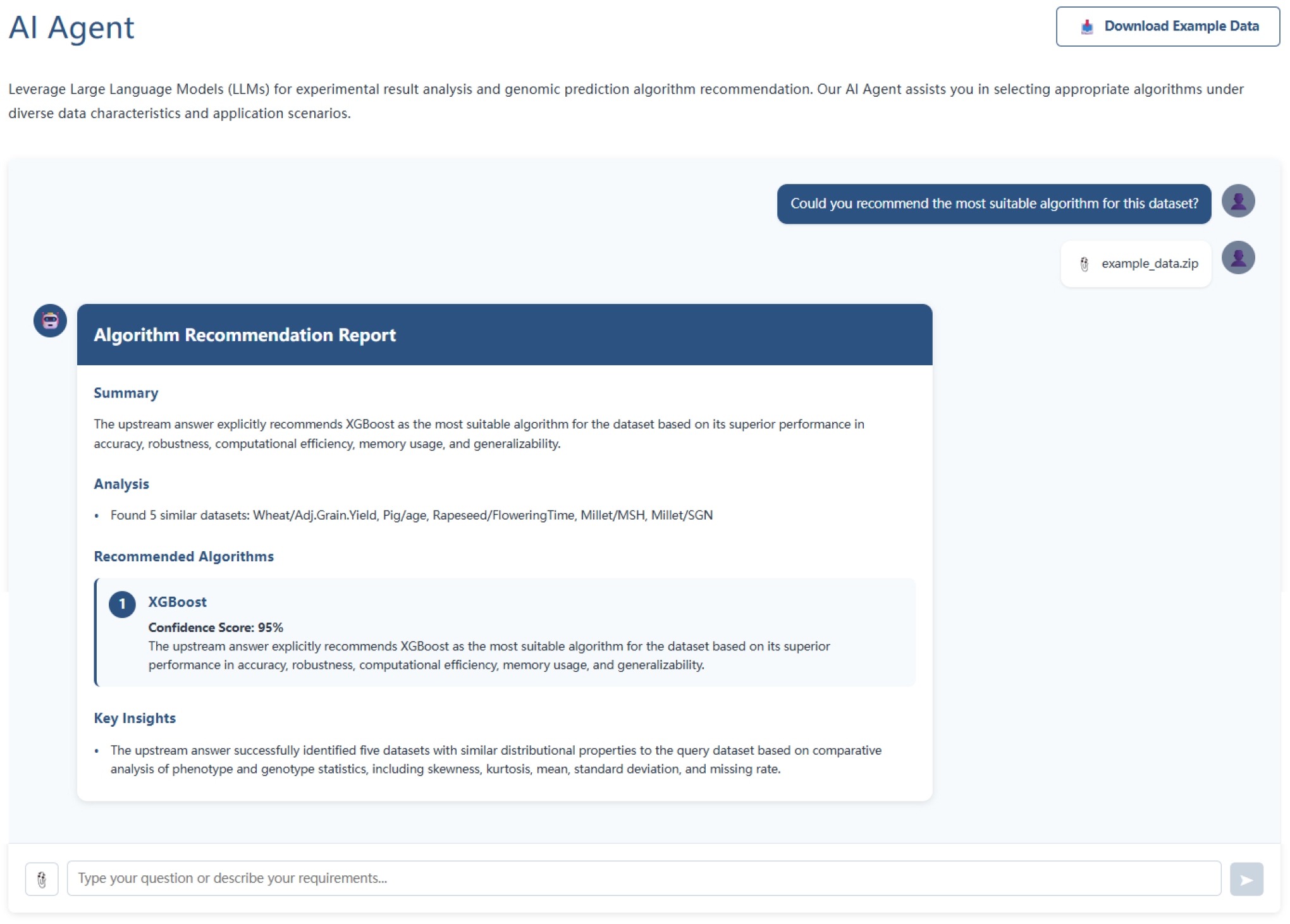
shows the user and agent chat process in algorithm recommendation tasks.

## Footnotes

1 https://huggingface.co/datasets/xw97/GPBench

2 https://github.com/xwzhang2118/GPBench

